# Improving cell-free expression of membrane proteins by tuning ribosome co-translational membrane association and nascent chain aggregation

**DOI:** 10.1101/2023.02.10.527944

**Authors:** Jan Steinküher, Justin A. Peruzzi, Antje Krüger, Miranda L. Jacobs, Michael C. Jewett, Neha P. Kamat

## Abstract

Cell-free gene expression (CFE) systems are powerful tools for transcribing and translating genes outside of a living cell. Given their diverse roles in nature, synthesis of membrane proteins is of particular interest, but their yield in CFE is substantially lower than for soluble protein. In this paper, we study factors that affect the cell-free synthesis of membrane proteins and develop a quantitative kinetic model of their production. We identify that stalling of membrane protein translation on the ribosome is a strong predictor of membrane protein synthesis and creates a negative feedback loop in which stalled peptide sequences quench ribosome activity through aggregation between the ribosome nascent chains. Synthesis can be improved by the addition of lipid membranes which incorporate protein nascent chains and, therefore, kinetically competes with aggregation. Using both quantitative modeling and experiment, we show that the balance between peptide-membrane association and peptide aggregation rates determines the total yield of synthesized membrane protein. We then demonstrate that this balance can be shifted by altering membrane composition or the protein N-terminal domain sequence. Based on these findings, we define a membrane protein expression score that can be used to rationalize the engineering of N-terminal domain sequences both of a native and computationally designed membrane proteins produced through CFE.

## Introduction

Cell-free gene expression (CFE) systems leverage cellular machinery to transcribe and translate genes outside of a living cell[1], [2]. Over the last two decades, CFE systems have grown from a molecular biology tool to a powerful shelf-stable and scalable biomanufacturing platform[3]–[7]. CFE systems have now been used to create wearable biosensors,[8] prototype metabolic pathways,[9], [10] rapidly screen drug candidates,[11], [12] and produce vaccines at the point of care[13], [14]. Thus, efforts to expand the capabilities of CFE systems could have a large impact on sustainable biomanufacturing, point-of-use biosensing, and therapeutic production.

One area that has posed a challenge with CFE systems has been the robust expression of membrane proteins. This is because membrane proteins require amphiphilic scaffolds to integrate into, similar to their synthesis in living cells. Because membrane proteins perform critical cellular functions in sensing, signaling, and energy regeneration, their inclusion in CFE systems is critical to expand the sensing and biomanufacturing capabilities of CFE systems.

Towards addressing this need, membranes and membrane mimetics have been included in CFE systems to integrate and improve expression of membrane proteins [15]. Inverted vesicles, formed from cellular membranes during extract preparation, have been used to retain membrane-associated functionality in CFE systems[6], [16]. However, the production of native vesicles requires overexpression of membrane components prior to lysis and is limited by the challenges associated with heterologous membrane protein production and furthermore, does not allow for tuning of membrane biophysical features, which may affect the final activity of an expressed membrane protein[17]. The ability to directly express membrane proteins into a membrane mimetic in a CFE system could circumvent these challenges. Synthetic membranes, in the form of liposomes and nanodiscs, have been used in CFE systems to improve expression of membrane proteins[17]–[20]. Experiments have established that membrane composition, available membrane area, and formation of co-translational membrane-bound ribosome complexes are crucial for successful CFE of membrane proteins[18], [21], [22]. However, each of the properties must be tuned to effectively produce properly folded, functional membrane proteins, limiting the ready adoption of membrane proteins into CFE systems[17], [22], [23]. The ability to predict optimal reaction conditions for CFE of membrane proteins could enable efficient membrane protein expression and consequently the rapid expansion of membrane functionality within cell-free systems.

To systematically improve CFE, insight from mechanistic models has proven useful for soluble protein expression. Recently, coarse-grained and multi-parameter models were used to quantitatively describe CFE systems, including sequence-specific predictions of translation and translation (TX/TL) kinetics[24]–[30]. However, to the best of our knowledge, a similar quantitative model for cell-free membrane protein synthesis does not exist. In this paper, we (i) compare CFE of a soluble and membrane protein and (ii) develop a quantitative model to describe cell-free membrane protein synthesis. We then (iii) apply this model to improve expression of a native and a computationally designed membrane protein by up to 50%.

## Results

### Expression of membrane proteins reduces the capacity of CFE systems to produce proteins

As a first step towards deconstructing the expression of membrane proteins in cell-free systems, we quantified CFE activity when expressing a model soluble protein, super folder green fluorescent protein (sfGFP), and a model membrane protein, mechanosensitive channel of large conductance (MscL) fused to monomeric enhanced green fluorescent protein (MscL-GFP)[23]. We produced proteins from plasmids encoding these proteins using the PURE system, which is based on 31 purified macromolecular components of the cellular transcription and translation machinery plus material and energy resources [31]. Fluorescent protein expression and folding can be monitored by measuring the resulting fluorescent signal as the cell-free reaction proceeds over time (**Fig. 1a**). We assumed that GFP fluorescence is a measurement of correctly folded protein yield. Apart from the overall fluorescence yield, two other quantities can be extracted from such experiments: the maximum synthesis rate and lifetime of the reaction. We compared these quantities for soluble protein, sfGFP, and membrane protein, MscL-GFP (blue and open black points **Fig. 1b**). We found that MscL-GFP yield, synthesis rate, and reaction timescale was two to three orders of magnitude lower than that of soluble sfGFP. Addition of lipid material in the form of 100 nm liposomes composed of 1,2-Dioleoyl-sn-Glycero-3-Phosphocholine (DOPC) enhanced some of the overall yield and synthesis rate (filled black points **Fig. 1b**). However, even in the presence of lipid vesicles, the reactions never came close to the performance of the cell-free system expressing soluble protein. Such large differences cannot be explained by differences of protein molecular weight (MW) or differences in the GFP variants. Instead, it appeared as though expression of MscL quenched the CFE reaction and therefore did not use the available resources efficiently. We wanted to understand this effect in more detail.

**Figure 1.**
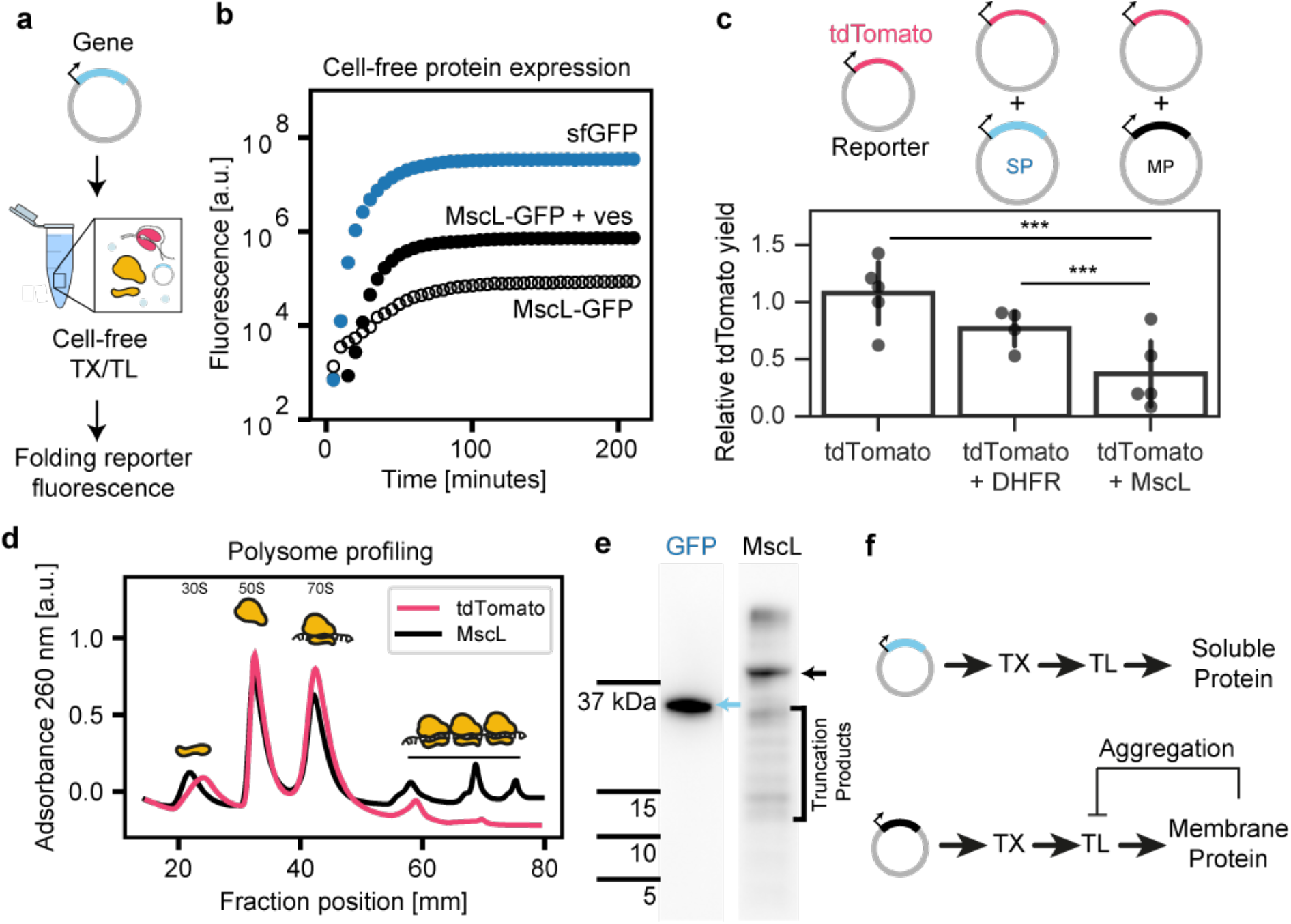
Expression of membrane protein quenches a cell-free expression (CFE) reaction. a) Schematic of the cell-free system and fluorescence assay for protein expression. b) Representative results for protein expression upon addition of plasmids coding soluble super-folder green fluorescent protein (sfGFP) or the mechanosensitive channel of large conductance-enhanced GFP fusion (MscL-GFP) with (+ ves) or without addition of 100 mM liposomes. c) Quenching of CFE is monitored by co-expressing fluorescent tdTomato with a soluble non-fluorescent dihydrofolate reductase (DHFR) (soluble protein, SP) or a non-fluorescent MscL (membrane protein, MP), respectively and normalized by maximum fluorescence of tdTomato alone. d) Polysome formation was detected (indicated by horizontal line) using a sucrose density gradient light (260 nm) adsorption profile after expression of tdTomato or MscL (no liposomes added). e) Western blot against the N terminus of soluble mEGFP and MscL-GFP demonstrates that more truncation products are produced during the expression of MscL-GFP. Black arrow indicates full length MscL-GFP. f) Proposed feedback loop that quenches translation by ribosome stalling/aggregation when expressing membrane protein.

To investigate the extent to which membrane protein expression inhibits cell-free expression reactions, we monitored the expression of a soluble, fluorescent protein, tdTomato, in the presence of a co-expressed soluble or membrane protein (**Fig. 1c**). Specifically, we compared the expression of fluorescent tdTomato (MW 54 kDa) with either soluble non-fluorescent dihydrofolate reductase (DHFR) (MW 24 kDa) or non-fluorescent MscL (MW 14 kDa) to tdTomato alone. In all three experiments there were no lipids added to the reaction. We found that co-expression of the soluble dihydrofolate reductase protein slightly decreased tdTomato yields, while co-expression of MscL decreased tdTomato yields by about 70 % relative to tdTomato expression alone. The capacity of membrane protein co-expression to significantly reduce expression of a soluble protein suggests that the expression of membrane proteins quenches CFE activity.

We wondered if the observed quenched CFE activity in the presence of co-expressed MscL is caused by aggregation of misfolded MscL peptides. We reasoned that MscL, which has large hydrophobic peptide segments, should demonstrate a higher propensity of misfolding in the absence of a lipid membrane to insert and fold into. As such, MscL peptides might aggregate during translation, stall the ribosome, and reduce the pool of ribosomes available for translation relative to soluble proteins. Accordingly, we measured ribosome aggregation and peptide fragmentation in our cell-free reactions as a function of expression of MscL or tdTomato. To measure ribosome aggregation, cell-free reactions were quenched on ice and transferred to a sucrose gradient. After ultracentrifugation, differently sized ribosome complexes sediment along the gradient. The presence of RNA material along the sucrose gradient was then measured by light adsorption. When soluble tdTomato was expressed, we obtained the characteristic peaks of 30S, 50S and 70S, corresponding to the small and large ribosomal subunits and assembled 70S ribosome (magenta trace **Fig. 1d**)[32], [33]. When MscL was expressed, we observed additional peaks that are assigned to polysomes, i.e., multiple ribosomes stalled along an RNA strand (black trace **Fig. 1d**). Previously, polysome formation was shown to be enhanced by attractive interactions between nascent chain complexes[34]. Because large attractive interactions should be present between the hydrophobic membrane protein residues, the increased interaction of membrane protein nascent chains would be expected to increase polysome formation as we observed. In addition to polysome formation, the stalling of ribosome complexes should give rise to incomplete protein products. Indeed, by probing the N-terminus of soluble GFP and MscL-GFP via western blot, we observed the presence of truncated protein products only when the membrane protein, MscL was expressed in contrast to when GFP was expressed (**Fig 1e, S1**). In summary, our results suggest a negative feedback loop by which expression of a membrane protein stalls, or quenches, ribosome activity due to aggregation of the ribosome-bound nascent chains in the PURE cell-free system (**Fig. 1f**).

### Kinetic model reveals balance between membrane association and aggregation

We next developed a kinetic model to describe different ribosome states during protein expression. We coarse-grained the cell-free synthesis into a series of elementary reactions (see Methods). We modeled the synthesis of mRNA as a first order reaction with rate constant *k_RNA_*. Translation initiation occurs by a second order reaction between free ribosome and mRNA with a rate *k_init_*. Ribosomes are deactivated with a rate constant *k_deg_*, a parameter which captures resource and energy depletion of the PURE system[26], [27]. This degradation is what leads to finite activity of the PURE system in previous models (the plateau in **Fig 1b**). The protein is synthesized with the previously determined rate constant *k_syn_* which leads to a growing nascent chain until the protein is fully synthesized and the ribosome is released from the mRNA. Finally, we modeled the effect of a blocked ribosomal binding site between individual ribosomes by not allowing binding of a ribosome to an occupied mRNA initiation site.

Compared to previous studies, which only considered ribosomes in solution, our model takes two additional states into account. First, we included a bound complex between the ribosome’s nascent chain and the membrane, which forms between the translated peptide and the membrane surface with binding/unbinding rate constants *k_+_* and *k_−_*. Second, we considered an aggregated, dysfunctional state which removes ribosomes from the system with a second order rate constant *k_agg_* due to encounters between RNA-bound ribosomes. In the latter case of aggregation, truncated protein products, with lengths of the aggerated nascent chain, are produced. We assumed that binding of the translated peptide to the membrane is reversible and is a function of the length of the synthesized protein. Specifically, we sought to capture the fact that a short, N-terminal segment of a not fully synthesized membrane protein will exhibit lower membrane affinity than a longer translated transmembrane segment of the same membrane protein. To accomplish this effect, we consider *k_+_* (binding) as constant and *k_−_* (unbinding) decaying exponentially with protein length *L*, an assumption we will revisit later. Fully synthesized protein folds with a rate *k_mat_*, corresponding to the GFP maturation rate, that we can compare to experimental values. This model was formalized as a set of elementary reactions (see Methods).

We proceeded to fit our model to experiments by comparing model-derived results to the fluorescence of the MscL-GFP folding reporter expressed in the presence of vesicles at different lipid concentrations. In our model, the vesicle concentration enters in the binding rate of ribosome-nascent chain complexes *M* to the membrane as 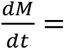 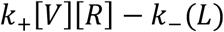 where *V* represents the concentration of vesicles and *R* is the free ribosome concentration. *V* is calculated from the vesicle size and total lipid concentration. The experiment reports the fluorescence of expressed MscL-GFP normalized to a baseline of McsL-GFP expression with zero lipid concentration *V* = 0. To fit the experimental data we also normalized the model output to GFP fluorescence at zero lipid concentration and assumed that all fully synthesized MscL-GFP molecules are fluorescent after the maturation time *k_mat_*. Considering the model simplifications and typical experimental error, the fit is satisfactory (**Fig. 2b**).

**Figure 2.**
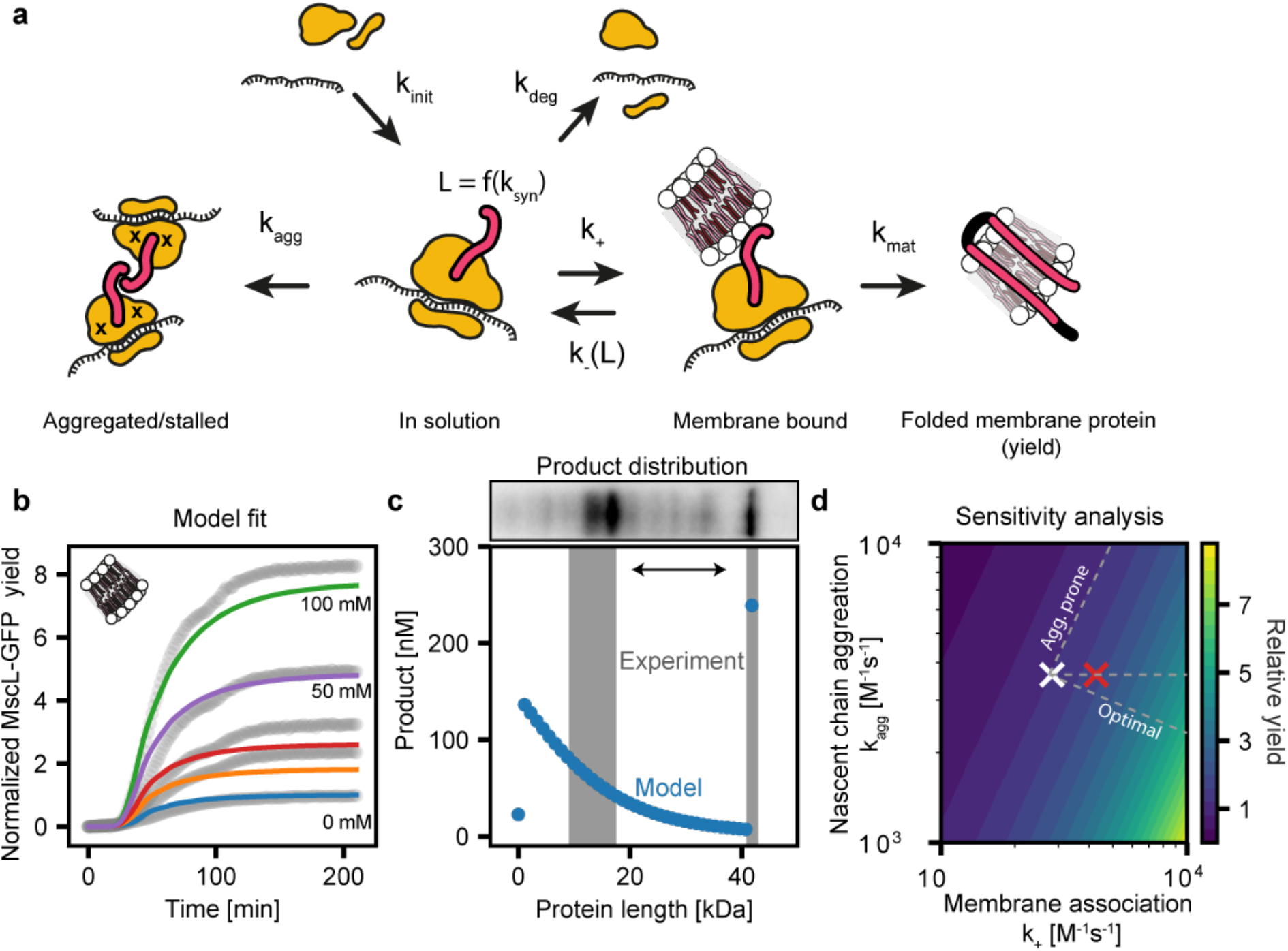
The balance of association of membrane protein nascent peptides to membranes or other ribosome-bound peptides determines the extent of quenching of a cell-free reaction. a) Overview of the kinetic model showing rate constants for initiation of translation *k_init_*, finite reaction lifetime due to resource depletion *k_deg_*, aggregation of nascent chains complex *k_agg_*, membrane association rate constant *k_+_* and *k_−_*(*L*), nascent chain length *L* that growths with rate *k_syn_* and *k_mat_* is the fluorescence reporter synthesis and maturation rate, respectively. b) Fit of the model to the data from Jacobs et al.[23] for MscL-GFP CFE with varying concentrations of liposome lipids (experimental data points are shown in grey and the best fit model is represented by colored lines) normalized to the zero lipid concentration. The experiments measure GFP fluorescence while the model calculates amount of full-length GFP, both quantities are referred to as “yield”. See main text for details. c) (Top) Gel image showing the distribution of protein products at the final time point of a CFE reaction expressing MscL-GFP in the presence of 10 mM DOPC liposomes. Molecular weight (MW) of aggregation and full-length bands approximately correspond to the MW defined by the x-axis below. Full-length MscL-GFP (46 kDa) is produced in addition to smaller truncation products. (Bottom) Model-derived truncation products are represented in blue while the gray columns indicate the regions of the gel with the highest detected protein density. Both experimental and model-derived data exhibit a gap in truncation product bands in the 20-40 kDa range, similar to the experimental result (western blot on top, arrow indicates the protein size range where less protein products are detected). d) Color map of full-length protein yield normalized to best fit values of *k_+_* and *k_agg_* (white cross). Increase of *k_+_* by hybrid membranes (red cross) and possible trajectories (dashed white lines) result in an increase of protein yield (“Optimal” and horizontal line) or no increase (“Aggregation prone”) .

We further established the validity of our model by comparing the calculated fitting parameters (Table 1) to literature values. Our model’s initiation rate lies between two previous estimates of 3 10^3^ M^−1^s^−1^ for the PUREfrex system by Doerr *et al*. and 175 10^6^ M^−1^s^−1^ for *in vitro* translation in optimized buffer conditions by Rudorf *et al*.[28], [35]. Similarly, our model’s first order transcription rate of a membrane protein, *k_RNA_*, is within error of the initial transcription rate in the PURE system (0.24 10^−9^ M^−1^s^−1^) obtained by measurements of RNA abundance in the PURE system by Gonzales *et al*. [27] (SI Note 1). In contrast, our model’s membrane association rate constant *k_+_* is one order of magnitude smaller than what was found using single molecule experiments between a peptide (GLP-1) and lipid membrane, with a binding rate of 1.0 10^4^ M^−1^s^−1^[36]. This difference might be due to differences in the peptide sequence between GLP1 and MscL. Additionally, binding of the nascent chain to the membrane requires correct alignment between a ribosomal exit tunnel and a membrane which reduces the binding rate compared to the free peptide GLP-1. While ribosomal association should impact membrane association, it would not be expected to contribute to unbinding of the nascent chain. Indeed, the unbinding rate *k_−_* aligns better with the experimental estimate for GLP-1 of 0.8 s^−1^. Combined, the fit of our model to previously generated data and its consistency with those found in literature demonstrates that our model adequately describes the experimental data by Jacobs *et al*.[18].

**Table 1.** Overview of model parameter and experimental values obtained from Gonzales *et al*. [27]. Standard deviation for fitted values is indicated by ± and sample size is n=7.

| Literature values |  |  | Fitted values |  |  |  |  |
| --- | --- | --- | --- | --- | --- | --- | --- |
| $k_{syn}$ [$s^{-1}$] | $k_{deg}$ [$10^{-6} s^{-1}$] | $[R]$ [$10^{-6} M$] | $k_{init}$ [$10^6 M^{-1}s^{-1}$] | $k_{RNA}$ [$10^{-9} Ms^{-1}$] | $k_+$ [$10^3 M^{-1}s^{-1}$] | $k_-$ [$s^{-1}$] | $k_{agg}$ [$10^3 M^{-1}s^{-1}$] |
| 0.33 | 385 | 2.4 | $5 \pm 3$ | $0.26 \pm 0.04$ | $2.8 \pm 0.2$ | $0.8 \pm 0.1$ | $3.6 \pm 0.4$ |

Next, we investigated the simulation trajectories in more detail. In the model, unbinding of ribosome-nascent chain complexes from the membrane becomes exponentially less likely when nascent chain sequences are longer than 10 amino acids. Thus, once the nascent chain is about 20 amino acids long, the ribosome-nascent chain complex does not unbind during the CFE lifetime. This means that ribosome-nascent chain complexes either aggregate early or integrate into a membrane co-translationally, where they are protected from aggregation. Competition between aggregation and integration results in a multimodal distribution of peptide products with a noticeable reduction or absence of products between unfinished, aggregated products of 10-20 kDa, and full-length protein of 40 kDa, both in the model calculations (**Fig. 2c**) and experiments (blot insert **Fig 2c**). These data indicate that the fate of each ribosome is determined early, when the synthesized protein is still rather short, by N-terminal domain binding affinity to the membrane and its aggregation propensity.

To gain quantitative insight into this effect, we varied the two rate constants *k_+_* and *k_agg_* in our simulation model (**Fig. 2d**). By calculating the yield of fully synthesized protein relative to the pair of the experimentally determined rates (*k_+_, k_agg_*), we found that excess aggregation quickly diminishes yield of full-length product, while a higher membrane association rate constant increases that yield (changes relative to white cross in **Fig. 2d**). These two rate constants (*k_+_* and *k_agg_*) can be tuned by changing the molecular components within the CFE reaction, such as the properties of the membrane and protein, allowing for increased protein yield. For example, changes in membrane composition might change *k_+_*, but should keep *k_agg_* constant (horizontal line in **Fig. 2d**). Recently, we used coarse grained simulations to show that hybrid polymer:lipid membranes can enhance peptide insertion rates by a factor of 1.5 relative to pure lipid membranes through a generic mechanism based on the generation of membrane packing defects.[37] Through this effect, association of the nascent-chain ribosome complex to the membrane will be enhanced by the same factor. By increasing *k_+_* in our model by 1.5 we predicted an increase in MscL yield (red cross **Fig. 2d**). The calculated 27 % yield increase is in almost exact agreement with the experimental result of 28 % ± 3 % improvement in MscL expression using the same hybrid polymer:lipid membranes. This agreement between the models and experimental data demonstrates the high predictive power of our modelling approach, as the polymer:lipid data was not used in the parameterization of the kinetic model.

Apart from changing the membrane composition, the protein sequence could also be altered to increase membrane protein yield. Our analysis predicts that an N-terminal domain sequence that both lowers aggregation propensity and increases membrane association would increase protein yield very effectively (“Optimal” trajectory in **Fig. 2d)**. However, sequences with high membrane affinity are often also prone to aggregation. Thus, there is a sequence space that will increase membrane association, but increase aggregation propensity to an even greater extent, leading to a constant or even reduction of yield (“aggregation prone” trajectory in **Fig. 2d**). Taken together, our modelling results suggest the need to balance membrane affinity and aggregation propensity of a membrane protein N-terminal domain to optimize membrane protein yield for a given membrane protein. Further, protein yield should be systematically enhanced by optimizing N-terminal domain membrane affinity.

### Diverse bacterial N-terminal domain sequences balance aggregation and membrane association

We wondered if we could design N-terminal peptide sequences in a way that promotes peptide affinity to synthetic membranes used in our study while limiting excessive peptide-peptide aggregation. For inspiration of suitable N-terminal sequences, we investigated naturally occurring sequences that have strong selection pressure against aggregation, which is generally toxic to cells. Initially, we again focused on MscL, which inserts co-translationally in bacteria, without assistance of the Sec translocon,[38], [39] similar to the cell-free expression system studied here. We hypothesized that as different bacterial species have large differences in their membrane composition, there might be a species with membrane compositions which better reflect the properties of simple synthetic membranes used in our cell-free reactions. Thus, by fusing a protein domain, which has evolved for optimal folding in membranes similar to our membrane mimetic, to the N-terminus of our protein, we might increase CFE protein yields. To identify such sequences, we calculated partitioning free energy Δ*G_wm_* values from water to a synthetic lipid bilayer interface for the first five residues from the N-termini from a diverse set of bacterial species identified by a consensus motif search (see Methods). Interestingly, the N-terminal helix of MscL is widely conserved between bacterial species, further promoting our investigation into the N-terminal domain sequence space (see Methods and Ref. [40]). The Δ*G_wm_* values correspond to partitioning free energies of peptide sequences from water to a zwitterionic phospholipid membranes interface, similar to the DOPC membranes added to the cell-free system[41]. We found a wide range of Δ*G_wm_* values between species. Notably, the native *E. coli* N-terminal domain sequence did not exhibit the highest membrane association energy. For our experiments we chose one sequence with comparable, larger, and smaller Δ*G_wm_* values (**Fig. 3a, b,c**). Additionally, we considered the N-terminal domain of another model membrane protein which expresses well in PURE cell-free reactions, LacY[42], [43]. Finally, we also investigated an artificial polyleucine sequence (LLLL), which would have the most favorable Δ*G_wm_* for membrane association. All five sequences were used to construct MscL chimeras between the different N-termini and the remaining *E. coli* MscL-GFP sequence. We then measured the cell-free reaction protein yields by GFP fluorescence. Between the naturally occurring sequences, we found a correlation (Pearson’s r=−0.9, p<0.05) with more favorable Δ*G_wm_* values increasing expression up to 40%, showing the potential in optimization of N-terminal domains for cell-free reactions. Interestingly, the synthetic polylysine construct LLLL fell outside of this correlation (yellow star in **Fig. 3c**), which is known to be very aggregation prone in solution[44]. The limited yield of protein expression observed for LLLL suggests this peptide promotes protein-protein aggregation, offsetting the large affinity to membrane association. In contrast, our results with the cell-derived peptide sequences suggest that nascent peptide sequences found in living organisms are selected against excess aggregation.

**Figure 3.**
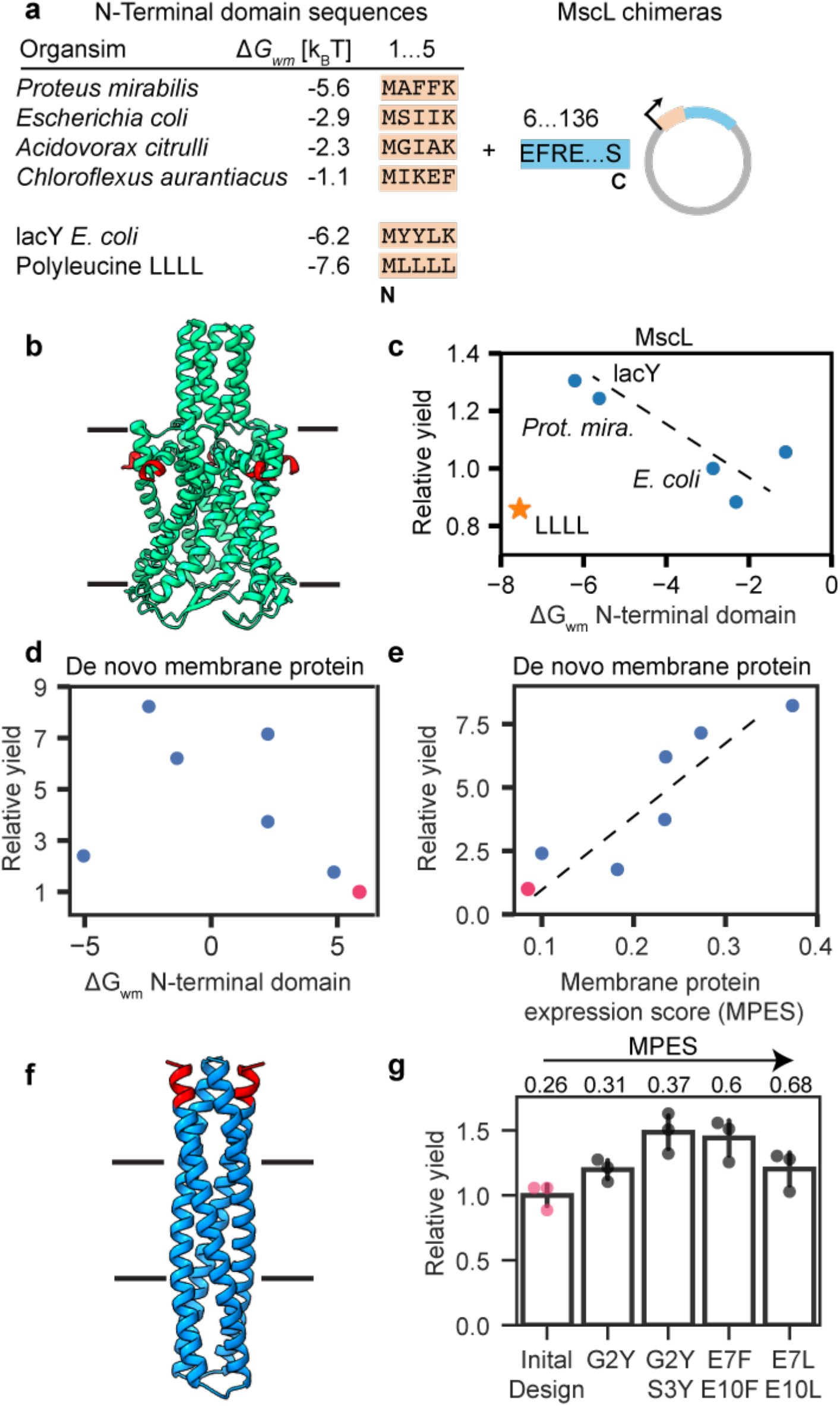
Rational engineering of membrane protein N-terminal domains for optimization of cell-free expression. a) Table of selected N-terminal domain sequences with Δ*G_wm_* (peptide partitioning free energy from water to lipid bilayer interface) values. The varying N-terminal domains (orange color) were fused to the N-terminus of MscL-GFP *(E. coli*, blue color) to obtain six different MscL-GFP chimeras. b) Cartoon of MscL inserted in the lipid membrane (black lines). Red indicates the N-terminal domain segment c) Resulting GFP fluorescence normalized to *E. coli* MscL (relative yield) of the six MscL-GFP chimeras plotted against the calculated Δ*G_wm_* values. d) Calculated Δ*G_wm_* values vs. GFP expression determined by fluorescence normalized to the worst expressing *de novo*-designed protein (red dots) show only weak correlation. e) Same relative GFP expression data as in panel d plotted against the membrane protein expression (MPES) score defined in the main text show a strong correlation (Pearson’s r=−0.9, p<0.05). f) Cartoon of de novo protein inserted in the lipid membrane (black lines). Red indicates N-terminal domain segment g) Improvement of de novo membrane protein yield by mutations that increase the MPES. MPES score is normalized between 0 and 1, where 1 indicates highest membrane affinity with lowest aggregation propensity. Numbers above bars show calculated MPES score. Each datapoint represents an independent experiment.

### Restoring the balance between membrane association and aggregation in n-termini of de-novo designed proteins

The excess aggregation of the synthetic polylysine construct made us curious to see if our insights could be used to improve expression of *de novo* membrane proteins, where aggregation often is a limiting factor for new designs. In a previous study, seven different *de novo* proteins with varying transmembrane domains were synthesized in the PURE system[45]. The design process resulted in transmembrane proteins not only with varying transmembrane lengths but also different N-terminal sequences, giving us the opportunity to test the effect of balancing aggregation and membrane affinity. Expression and protein yield, as measured by GFP fluorescence, between the seven designs varied about eight-fold (**Fig. 3d**). Comparison between relative expression and Δ*G_wm_* values reveals only a weak correlation, further strengthening our assumption that synthetic N-terminal domain sequences lack the necessary balance between membrane association and aggregation propensity. To assess aggregation propensity, we considered the CamSol solubility score, which is based on a phenomenological amino acid aggregation scale [46]. We scale both Δ*G_wm_* and CamSol solubility score between 1 and 10, where 1 is largest Δ*G_wm_* value and lowest CamSol score. This analysis gives two values, s_1_ (Δ*G_wm_*) and s_2_ (CamSol), between 1 and 10. We define the membrane protein expression score (MPES) as s_1_·s_2_/10 where 10 would predict best expression. The correlation between measured expression and score is highly significant (Pearson’s r = 0.9, p < 0.01), showing that synthetic sequences need to consider both aggregation propensity and membrane affinity (**Fig. 3e**). Similarly, the MPES correctly predicted the polyleucine sequence scoring lowest, while the best expression construct scored highest among the MscL chimeras (**Fig. S2**). Motivated by these results we asked if we could add mutations that increase MPES to improve expression of otherwise low expressing membrane protein. We considered the worst expressing construct, the 20 Å thick transmembrane protein, and generated a single point mutant G2Y, and double mutants (G2Y, S3Y), (E7F, E10F) and (E7L, E10L) which increase MPES (**Fig 3f, g**). As predicted all five constructs improved expression, with the best expression by mutant (G2Y, S3Y) which improved expression yield by approximately 50%. Together, these results demonstrate cell-free expression of membrane proteins can be improved by systematically changing the N-terminal domain sequence and that the MPES score provides a metric to guide sequence design.

## Discussion

Here, we have taken a closer look at biophysical features of cell-free membrane protein expression. By characterizing polysome formation and truncation of protein products for soluble and membrane proteins, we established that the aggregation state of protein expressed in the PURE system changes with membrane protein expression. Our coarse-grained model emphasizes that increases in membrane protein yield with the addition of a membrane surface is a kinetic effect, and not due to saturation of available membrane surface. It also shows that one cannot simply consider CFE of membrane proteins as an equilibrium between membrane-bound and solution-dispersed protein. Instead, the total yield of protein is strongly influenced by a negative feedback loop leading to ribosome aggregation and stalling. An important result of our study is the competition between membrane association rate k+[*V*][*R*] and aggregation rate *k_agg_*[*R*]^2^. At fixed initial ribosome [*R*] and vesicle concentration [*V*], the balance is determined by the ratio of the rate constants *k_+_* and *k_agg_*. Our results are corroborated by experimental results from other groups who found that membrane protein yield is reduced by deletion of the N-terminal domain, but increased by anchoring of the nascent chain by NTA-His complexes onto the membrane surface[39], [47]. Anchoring of expressed proteins to the membrane will increase *k_+_* while deletion of the N-terminal domain will expose the very hydrophobic transmembrane segment to the solute, increasing *k_agg_*. Additionally, Eaglesfield *et al*. have shown that excess aggregation by deletion of the N-terminal domain can be improved by artificial ribosome anchoring to the membrane, fully consistent with our model[39]. Our results are also consistent with results from Harris *et al*. that have shown large increases in yield of both aggregated and membrane inserted protein with changes in membrane composition[22], a result that cannot be explained by equilibrium partitioning models but corroborates our described feedback mechanism.

To improve membrane protein yield, we examined how the rate constant *k_+_* could be altered to change membrane association. The rate constant *k_+_* can be tuned by the molecular properties of the lipid membrane, an effect we have previously studied quantitively, demonstrating peptide insertion rates into polymer:lipid membranes are increased by generation of lipid packing defects[37]. Further effects on *k_+_* might be expected by other changes in membrane composition, e.g., headgroup charge or hydrocarbon chain saturation[42]. Less is known about the effects of membrane composition on the dissociation rate constant *k_−_*. Recently it was shown that CFE membrane protein yield can be improved by increasing membrane viscosity[17]. Membrane viscosity might lower *k_−_* as dynamics of protein adsorbed to the interface will be slower. Lower *k_−_* values should lead to larger membrane affinity and, in this way, could improve yield. To understand these effects quantitively would be useful to obtain more systematic data on peptide or nascent chain unbinding kinetics with membrane composition.

Apart from the contribution of membrane composition, both *k_+_* and *k_−_* will depend on the protein sequence. Unfortunately, no systematic prediction between these rate constants and sequence exists. In principle molecular dynamic simulation or single molecule experiments could provide these rates. Instead of relying on these low-throughput methods, we utilized a result from our model that a protein’s N-terminal domain will determine success or aggregation early on during protein synthesis. This motivated us, even if the cell-free system is clearly an out of equilibrium system, to approximate this binding step as an equilibrium between the nascent chain complex and membrane surface with a partitioning free energy of Δ*G_wm_*. We justify this approximation because the short nascent chain binds and unbinds faster from the membrane than the nascent chain elongation, which provides the system time to sample its equilibrium distribution. If aggregation is avoided, more favorable N-terminal domain Δ*G_wm_* values would always predict higher protein yield, which we indeed observed using MscL-chimeras. These results suggest that evolved sequences from biological organisms have balanced membrane affinity with aggregation propensity. As expected by this reasoning, a synthetic polylysine sequence, which should have both high membrane affinity and a high propensity to aggregate, did not improve expression as the most favorable Δ*G_wm_* value might suggest. Based on our modelling results, we defined the MPES scale, which allows us to quantify the balance of aggregation vs membrane association by previously determined empirical scales. We believe that our approach, which combines mechanistic insight with rational engineering, could be applied to increase the expression of a wide array of membrane proteins.

### Limitations and possible extensions of our model

Our model made a series of simplifications. For example, translation initiation can be more explicitly modeled as a multi-step process, or codon-specific elongation rates might be considered[28], [35]. We would expect that by these additions, the predications for truncated products would become sharper and better resolve the band structure of the western blots. To account for sequence specific variations of *k_RNA_* and *k_init_* our model could be combined with approaches that consider thermodynamics of ribosome-RNA binding or transcription initiation rates[24], [25].

Importantly, the main result of the competition between membrane association and aggregation is robust against variations in these parameters. We suggest that the crudest simplification is that protein of any length at the membrane is protected from aggregation. Instead of the simplified picture of protein maturation with a single rate constant *k_mat_*, proteins on the membrane surface insert and fold in a multi-step process. For example, force pulling experiments suggest that multi-pass protein fold by insertion of harpins, divided by barriers on the order of ~ 10 k_B_T[48]. If the lifetimes of unfolded protein on the membrane surface are comparable, the frequency of collisions between two not-yet folded proteins on the same or different liposomes leads to additional aggregation pathways (**Fig. 4**). In other words, depending on the stability of the folding pathway, co-translational insertion does not fully protect membrane proteins from aggregation as they might interact via intra- or inter-membrane interactions during the folding process. This mode of interaction provides a possible explanation for the weaker bands of truncated product in **Fig. 2c** which are not captured by the current model. Consistent with these ideas, a seven-pass Rhodopsin was observed to express with a different dependence on membrane composition than the simpler, two-pass MscL studied here and previously[18]. By additional measurements these short comings can be addressed systematically. In Peruzzi *et al*. the stability of hydrophobically mismatched *de novo* proteins was studied in detail using experimental and computational approaches[45]. This study showed that small changes to membrane protein stability by membrane deformation indeed systemically change protein yield, consistent with the additional aggregation pathways discussed here. Hydrophobic mismatch is straightforward to include quantitatively in our model but would require separate measurements of membrane protein association and insertion kinetics. Even larger changes of the of rate constants *k_f_* and *k_r_* with membrane composition, maybe even favoring the unfolded state, would clearly lower protein yield independent of the here described feedback effects.

**Figure 4.**
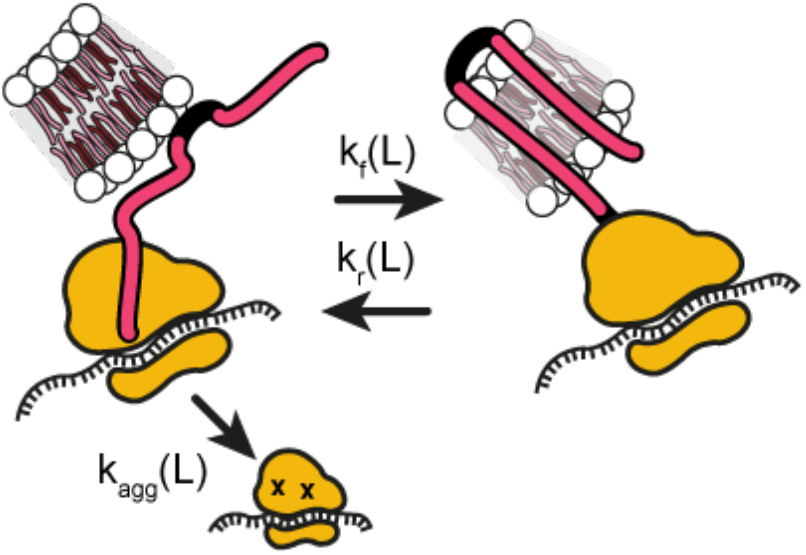
Possible addition considering stability of the membrane folding process at the membrane against aggregation. The protein folding and unfolding pathway, represented by the rates *k_f_* and *k_r_*, might depend on the on the protein length *L*. If these rates are in the same order as protein-protein collision rates, additional aggregation pathways appear.

Experimentally we have studied the PURE system. We speculate that our main results hold in principle for other CFE systems and maybe even for primitive cells. Clearly, in different CFE systems additional molecular players might become important. A possible modification of our model would be to allow for additional states that protect from aggregation. For example, chaperones might be present in crude cell extracts. To study such processes in detail it would be interesting to spike the PURE system with chaperones, or similar molecules, and observe their ability to effectively suppress aggregation and thereby improve membrane protein yield.

### Conclusion

We have developed a kinetic model for cell-free membrane protein expression. Our results provide a unified framework to understand CFE of membrane proteins and are consistent with a large number of experimental data. In principle, our model might be adjusted for each membrane protein, e.g., by parametrization of sequence-specific aggregation and membrane association rate constants. This approach is limited by its low throughput. Instead, we have used the developed intuition to define a membrane protein expression score that can be readily calculated and demonstrate the benefit that biophysical insight can provide on engineering bottom-up synthetic biology systems.

## Material and Methods

### Used plasmids and templates

MscL-GFP, sfGFP, and tdTomato were prepared as described previously[13], [23]. DNA templates for experiments performed in Figure 3, including chimeric MscL and de-novo designed proteins, were ordered as gBlocks from Twist Biosciences with a T7 promoter and terminator, as well as a ribosome binding site (**Table S1**). gBlocks were amplified via PCR and purified using a PureLink PCR Purification Kit (Invitrogen). PCR products were used directly in cell-free reactions.[31]

### Assembly of cell-free expression reaction and fluorescence readout

Protein expression was performed with the PURExpress In Vitro Protein Synthesis kit (E6800, NEB) according to the manufacturer’s instructions. 30 μL reactions were assembled with a final concentration of 10 mM of lipid and 3.3 nM plasmid (or approximately 200 ng). Reactions were allowed to progress at 37 °C for 3 hours. All fluorescence experiments were performed on a plate reader (Molecular Devices Spectra Max i3) at 37 °C with a reaction volume of 30 μL. Fluorescence of sfGFP and transmembrane domain-GFP fusion proteins (MscL and *de novo*-designed proteins) was measured with an excitation of 480 nm and emission at 507 nm. tdTomato was excited at 553 nm and emission collected at 581 nm.

### Sucrose gradient experiments

Sucrose gradients were prepared from gradient buffer using a Biocomp Gradient Master as described previously[32]. PURE cell-free reactions of 30 μL with plasmid coding for MscL or tdTomato were prepared and incubated at 37°C for 1.5 hours. Reactions were quenched by putting them on ice. Sucrose gradients were prepared from gradient buffer (20 mM Tris–HCl (pH 7.5 at 4 °C), 100 mM NH_4_Cl, 10 mM MgCl_2_) with 10 and 40% sucrose in SW41 polyclear centrifuge tubes (Seton Scientific) using a Biocomp Gradient Master and chilled to 4 °C. Cell-free reactions were diluted with 200 μl gradient buffer and layered onto chilled gradients. The gradients were ultra-centrifuged at 41,000 rpm for 3 hours at 4°C (Optima L-80 XP ultracentrifuge (Beckman-Coulter)). Gradients were analyzed with a Piston Gradient Fractionator™ (Biocomp) coupled to a Triax^TM^ FC-2 UV-260/280 flow cell. Traces of 260 nm light adsorption versus elution volumes were obtained for each gradient and adjusted by a blank sucrose sample.

### Western blots

Cell-free expressed proteins were analyzed by western blot to observe the presence of truncation products. Cell-free expressed protein samples were run on a 16.5% Tricine Mini-PROTEAN Precast Protein Gel to enhance the separation of smaller protein products. Wet transfer was performed onto a PVDF membrane (Bio-Rad) for 45 min at 100 V. Membranes were then blocked for an hour at room temperature in 5% milk in TBST (pH 7.6: 50 mM Tris, 150 mM NaCl, HCl to pH 7.6, 0.1% Tween) and incubated for 1 hour at room temperature or overnight at 4 °C with primary solution (anti-Flag, diluted 1:1000 in 5% milk in TBST). Primary antibody solution was decanted, and the membrane was washed three times for 5 minutes in TBST and subsequently incubated in secondary solution at room temperature for 1 hour (HRP-anti-Mouse (CST 7076) diluted 1:3000 in 5% milk in TBST). Membranes were then washed in TBST, incubated with Clarity Western ECL Substrate (Bio-Rad) for 5 min, and imaged in an Azure Biosystems c280 imager. Uncropped western blots are shown in Figure S1.

### Motif search

Annotated MscL homologs were downloaded from UniProt. N-terminal domains between 9 and 20 amino acids were identified by their annotation and extracted, yielding 181 sequences. These sequences were uploaded to the motif discovery tool MEME (v. 5.4.1)[49]. The multilevel consensus sequence MSIIKEFR appeared in 43 N-terminal domains, for which we calculated partitioning free energies as described below.

### Partitioning free energy and solubility score calculation

Partitioning free energies were calculated using MPEx (v 3.3.0) using the “Interfacial scale” with 100% Helicity with the setting “No End Groups” at 0 mV bilayer surface charge[50]. Solubility was calculated by the CamSol webserver (https://www-cohsoftware.ch.cam.ac.uk/index.php) using the “CamSol Intrinsic” setting.

### Model reactions and fit to data

RNA synthesis

1. 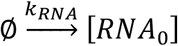

Ribosome binding RNA + synthesis in solution

2. 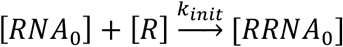

3. 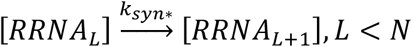

4. 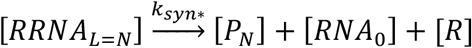

5. 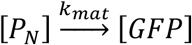

Ribosome binding / unbinding vesicle

6. 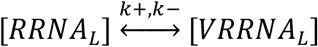

Co-translational protein synthesis and membrane protein maturation

7. 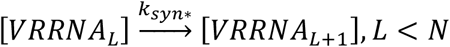

8. 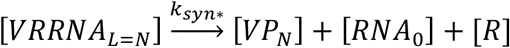

9. 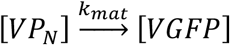

Ribosome degradation / fall off

10. 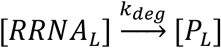

Ribosome aggregation

11. 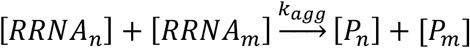

We considered the elementary reactions for synthesis of MscL-GFP with a total length of 1170 nucleotides. For computational efficiency the synthesis was modelled in steps of 10 amino acids, meaning that 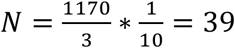 with a scaled synthesis rate 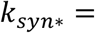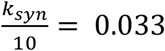 amino acids/s. For a nascent chain of length *L* the unbinding rate constant *k_−_* was scaled by exp (−*L* * 10) to reflect the incrase of membrane affinity with protein length. We account for proteins synthesized in solution [GFP] and at the membrane surface [VGFP]. For comparison with experiments, we calculated the total yield of [GFP]+[VGFP]. For Figure 2c the truncated protein of length *L* [*P*_L_] was reported.

The model reactions were implemented in python (v. 3.7.4) using the gillespy2 package (v. 1.6.7) by numerical integration of the ordinary differential equations defined above. The fit was performed by minimizing the sum of squared differences between model total GFP yield and experimental trajectories for all vesicle concentrations (global fit) using gp_minimize from scikit-optimize (v. 0.9). The fitting routine gp_minimize was performed by 1000 evaluation of the model function and otherwise default parameter values. Standard deviation of the fitted values was calculated from six gp_minimize runs. Further data processing was done using numpy (v. 1.21.4).

## Acknowledgments

This research was supported in part by the National Science Foundation under Grant No. 1844336 (N.P.K., M.C.J., J.S.) and No. 2145050 (N.P.K.), by the Army Contracting Command No. W911NF-22-2-0246 and W52P1J-21-9-3023 (N.P.K., M.C.J.) and through the computational resources and staff contributions provided for the Quest high performance computing facility at Northwestern University which is jointly supported by the Office of the Provost, the Office for Research, and Northwestern University Information Technology. J.A.P. gratefully acknowledges support from the Ryan Fellowship, the International Institute for Nanotechnology at Northwestern University, and an NSF Graduate Research Fellowship. J.S. acknowledges financial support from the Bundesministerium für Bildung und Forschung (BMBF) project NEUZELL (grant no. 031B1333).

## Notes

### Competing Interest Statement

The authors have declared no competing interest.

