## Supplementary Information for "Improving cell-free expression of membrane proteins by tuning ribosome co-translational membrane association and nascent chain aggregation"

### SI Note 1 – Initial transcription rate

Gonzales et al measured parameters for transcription in the PURE system modelled as Michaelis-Menten kinetics. We calculate the initial transcription velocity $\frac{dRNA}{dt} = \frac{k_r[DNA]}{K_r+[DNA]}$ with their parameter $k_r = 2894\ \frac{\text{nM}}{\text{h}}$ and $K_r = 3.67$ nM and the DNA concentration used in the experiments by Jacobs et al of 1.56 nM and found $\frac{dRNA}{dt} \approx 0.24\ \frac{\text{nM}}{\text{s}}$.

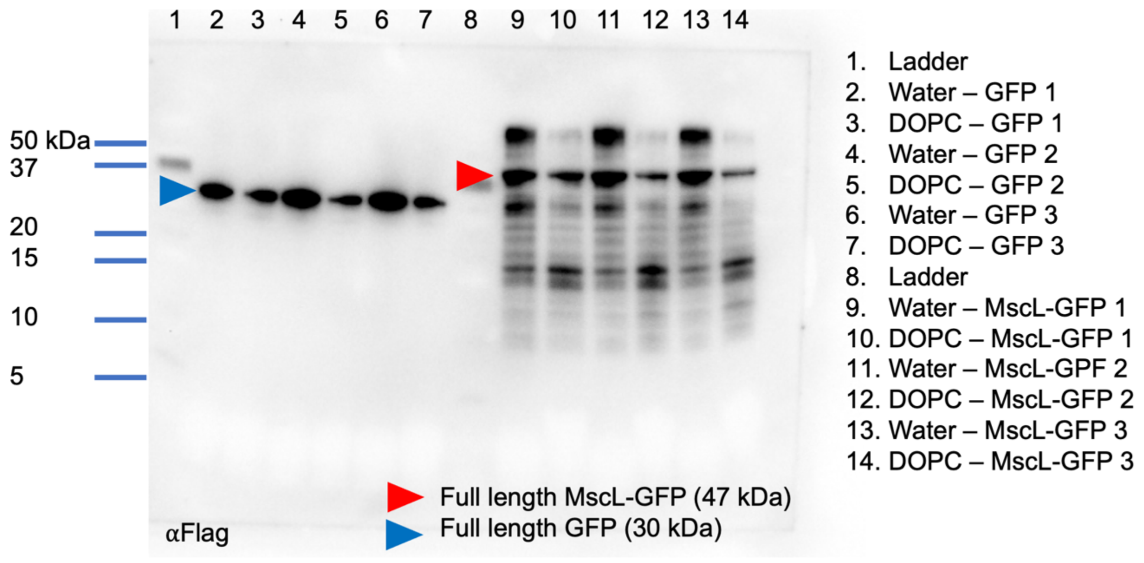


**Figure S1.** Uncropped western blot against the N-terminus of GFP and MscL-GFP with and without membranes. A Flag tag was added to the N-terminus of both proteins, enabling truncation products to be observed via Western blot. Minimal truncation was observed when expressing soluble GFP (Lanes 2-7), but truncation products were observed when MscL-GFP was expressed in the presence and absence of DOPC liposomes (10 mM lipid). Interestingly, more truncation products and higher molecular weight bands appear when MscL-GFP is expressed without liposomes. Experimental conditions were repeated 3 times (n=3) and observed on a single Western blot.

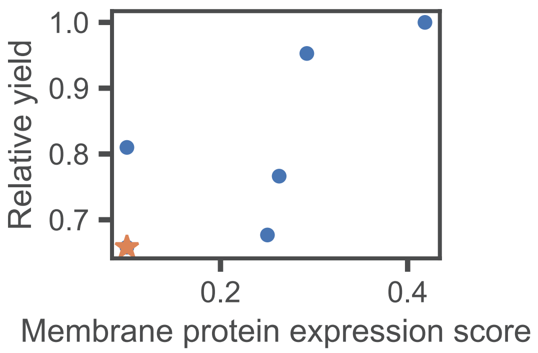


**Figure S2.** MPES score calculated for data from Fig. 3c. Polyleucine is indicated by a star.

**Table S1.** DNA sequences for constructs found in Figure 1 and Figure 2. All constructs were placed under the control of a T7 promoter.

| Construct | DNA Sequence |
|---|---|
| sfGFP | ATGAGCAAAGGTGAAGAACTGTTTACCGGCGTTGTGCCGAT<br>TCTGGTGGAACTGGATGGCGATGTGAACGGTCACAAATTCA<br>GCGTGCGTGGTGAAGGTGAAGGCGATGCCACGATTGGCAA<br>ACTGACGCTGAAATTTATCTGCACCACCGGCAAACTGCCGG<br>TGCCGTGGCCGACGCTGGTGACCACCCTGACCTATGGCGT<br>TCAGTGTTTTAGTCGCTATCCGGATCACATGAAACGTCACG<br>ATTTCTTTAAATCTGCAATGCCGGAAGGCTATGTGCAGGAA<br>CGTACGATTAGCTTTAAAGATGATGGCAAATATAAAACGCG<br>CGCCGTTGTGAAATTTGAAGGCGATACCCTGGTGAACCGC<br>ATTGAACTGAAAGGCACGGATTTTAAAGAAGATGGCAATAT<br>CCTGGGCCATAAACTGGAATACAACTTTAATAGCCATAATG<br>TTTATATTACGGCGGATAAACAGAAAAATGGCATCAAAGCG<br>AATTTTACCGTTCGCCATAACGTTGAAGATGGCAGTGTGCA<br>GCTGGCAGATCATTATCAGCAGAATACCCCGATTGGTGATG<br>GTCCGGTGCTGCTGCCGGATAATCATTATCTGAGCACGCA<br>GACCGTTCTGTCTAAAGATCCGAACGAAAAAGGCACGCGG<br>GACCACATGGTTCTGCACGAATATGTGAATGCGGCAGGTAT<br>TACGTGGAGCCATCCGCAGTTCGAAAAATAA |
| MscL-GFP | ATGGGCCATCATCATCATCATCATCATCATCATCACAGCAG<br>CGGCCATATCGACGACGACGACAAGCATATGAGCATTATTA<br>AAGAATTTCGCGAATTTGCGATGCGCGGGAACGTGGTGGA<br>TTTGGCGGTGGGTGTCATTATCGGTGCGGCATTCGGGAAG<br>ATTGTCTCTTCACTGGTTGCCGATATCATCATGCCTCCTCTG<br>GGCTTATTAATTGGCGGGATCGATTTTAAACAGTTTGCTGT<br>CACGCTACGCGATGCGCAGGGGGATATCCCTGCTGTTGTG<br>ATGCATTACGGTGTCTTCATTCAAAACGTCTTTGATTTTCTG<br>ATTGTGGCCTTTGCCATCTTTATGGCGATTAAGCTAATCAAC<br>AAACTGAATCGGAAAAAAGAAGAACCAGCAGCCGCACCTG<br>CACCAACTAAAGAAGAAGTATTACTGACAGAAATTCGTGATT<br>TGCTGAAAGAGCAGAATAACCGCTCTGGATCCGGCGGCGG<br>CAGCGAAAACCTGTATTTTCAGGGCATGGTGAGCAAGGGC<br>GAGGAGCTGTTCACCGGGGTGGTGCCCATCCTGGTCGAGC<br>TGGACGGCGACGTAAACGGCCACAAGTTCAGCGTGTCCGG<br>CGAGGGCGAGGGCGATGCCACCTACGGCAAGCTGACCCT<br>GAAGTTCATCTGCACCACCGGCAAGCTGCCCGTGCCCTGG<br>CCCACCCTCGTGACCACCCTGACCTACGGCGTGCAGTGCT<br>TCAGCCGCTACCCCGACCACATGAAGCAGCACGACTTCTT<br>CAAGTCCGCCATGCCCGAAGGCTACGTCCAGGAGCGCACC<br>ATCTTCTTCAAGGACGACGGCAACTACAAGACCCGCGCCG<br>AGGTGAAGTTCGAGGGCGACACCCTGGTGAACCGCATCGA<br>GCTGAAGGGCATCGACTTCAAGGAGGACGGCAACATCCTG<br>GGGCACAAGCTGGAGTACAACTACAACAGCCACAACGTCT<br>ATATCATGGCCGACAAGCAGAAGAACGGCATCAAGGTGAA<br>CTTCAAGATCCGCCACAACATCGAGGACGGCAGCGTGCAG<br>CTCGCCGACCACTACCAGCAGAACACCCCCATCGGCGACG<br>GCCCCGTGCTGCTGCCCGACAACCACTACCTGAGCACCCA<br>GTCCAAGCTGAGCAAAGACCCCAACGAGAAGCGCGATCAC |

| | |
|---|---|
| | ATGGTCCTGCTGGAGTTCGTGACCGCCGCCGGGATCACTC<br>TCGGCATGGACGAGCTGTACAAGTAA |
| tdTomato | ATGGTGAGCAAGGGCGAGGAGGTCATCAAAGAGTTCATGC<br>GCTTCAAGGTGCGCATGGAGGGCTCCATGAACGGCCACGA<br>GTTCGAGATCGAGGGCGAGGGCGAGGGCCGCCCCTACGA<br>GGGCACCCAGACCGCCAAGCTGAAGGTGACCAAGGGCGG<br>CCCCCTGCCCTTCGCCTGGGACATCCTGTCCCCCCAGTTC<br>ATGTACGGCTCCAAGGCGTACGTGAAGCACCCCGCCGACA<br>TCCCCGATTACAAGAAGCTGTCCTTCCCCGAGGGCTTCAAG<br>TGGGAGCGCGTGATGAACTTCGAGGACGGCGGTCTGGTGA<br>CCGTGACCCAGGACTCCTCCCTGCAGGACGGCACGCTGAT<br>CTACAAGGTGAAGATGCGCGGCACCAACTTCCCCCCCGAC<br>GGCCCCGTAATGCAGAAGAAGACCATGGGCTGGGAGGCCT<br>CCACCGAGCGCCTGTACCCCCGCGACGGCGTGCTGAAGG<br>GCGAGATCCACCAGGCCCTGAAGCTGAAGGACGGCGGCC<br>ACTACCTGGTGGAGTTCAAGACCATCTACATGGCCAAGAAG<br>CCCGTGCAACTGCCCGGCTACTACTACGTGGACACCAAGC<br>TGGACATCACCTCCCACAACGAGGACTACACCATCGTGGA<br>ACAGTACGAGCGCTCCGAGGGCCGCCACCACCTGTTCCTG<br>TACGGCATGGACGAGCTGTACAAGTAA |
| DHFR | ATGATCAGTCTGATTGCGGCGTTAGCGGTAGATCGCGTTAT<br>CGGCATGGAAAACGCCATGCCGTGGAACCTGCCTGCCGAT<br>CTCGCCTGGTTTAAACGCAACACCTTAAATAAACCCGTGAT<br>TATGGGCCGCCATACCTGGGAATCAATCGGTCGTCCGTTG<br>CCAGGACGCAAAAATATTATCCTCAGCAGTCAACCGGGTAC<br>GGACGATCGCGTAACGTGGGTGAAGTCGGTGGATGAAGCC<br>ATCGCGGCGTGTGGTGACGTACCAGAAATCATGGTGATTG<br>GCGGCGGTCGCGTTTATGAACAGTTCTTGCCAAAAGCGCA<br>AAAACTGTATCTGACGCATATCGACGCAGAAGTGGAAGGC<br>GACACCCATTTCCCGGATTACGAGCCGGATGACTGGGAAT<br>CGGTATTCAGCGAATTCCACGATGCTGATGCGCAGAACTCT<br>CACAGCTATTGCTTTGAGATTCTGGAGCGGCGGTAA |
| Flag-mEGFP | ATGGGTCATCATCACCACCATCACCATCATCATCACAGCAG<br>CGGTCATATCGATTATAAAGATGATGACGACAAACATATGG<br>TGAGCAAAGGCGAAGAACTGTTTACCGGTGTTGTTCCGATT<br>CTGGTTGAACTGGATGGTGACGTTAATGGTCACAAATTTTC<br>AGTTAGCGGTGAAGGCGAAGGTGATGCAACCTATGGTAAA<br>CTGACCCTGAAATTTATCTGTACCACCGGCAAACTGCCGGT<br>GCCGTGGCCGACACTGGTTACCACACTGACCTATGGTGTT<br>CAGTGTTTTAGCCGTTATCCGGATCACATGAAACAGCACGA<br>TTTCTTTAAAAGCGCAATGCCGGAAGGTTATGTTCAAGAAC<br>GTACCATCTTCTTTAAGGATGACGGCAACTATAAAACCCGT<br>GCCGAAGTTAAATTTGAAGGTGATACCCTGGTGAATCGCAT<br>TGAACTGAAAGGCATTGATTTCAAAGAGGATGGTAATATCC<br>TGGGCCACAAACTGGAATATAATTATAATAGCCACAACGTG<br>TACATCATGGCCGACAAACAGAAAAATGGCATCAAAGTGAA<br>CTTCAAGATCCGCCATAATATTGAAGATGGTTCAGTTCAGC<br>TGGCCGATCATTATCAGCAGAATACCCCGATCGGTGATGGT<br>CCTGTTCTGCTGCCTGATAATCATTATCTGAGCACCCAGAG<br>CAAACTGAGCAAAGATCCGAATGAAAAACGTGATCACATGG |

| | |
|---|---|
| | TCCTGCTGGAATTTGTTACCGCAGCAGGTATTACCTTAGGT<br>ATGGATGAACTGTACAAATAA |
| Flag-MscL-GFP | ATGGGTCATCATCACCACCATCACCATCATCATCACAGCAG<br>CGGTCATATCGATTATAAAGATGATGACGACAAACATATGA<br>GCATCATCAAAGAATTTCGCGAGTTTGCAATGCGTGGTAAT<br>GTTGTTGATCTGGCAGTTGGTGTTATTATTGGTGCAGCCTT<br>TGGTAAAATTGTTAGCAGCCTGGTTGCAGATATTATCATGC<br>CTCCGCTGGGTCTGCTGATTGGTGGTATTGATTTTAAACAG<br>TTTGCAGTGACCCTGCGTGATGCACAGGGTGATATTCCGG<br>CAGTTGTTATGCATTATGGTGTGTTTATTCAGAACGTGTTCG<br>ATTTTCTGATTGTGGCCTTTGCCATTTTCATGGCCATTAAAC<br>TGATCAACAAACTGAACCGCAAGAAAGAAGAACCGGCAGC<br>AGCACCGGCACCGACCAAAGAAGAAGTTCTGCTGACCGAA<br>ATTCGTGATCTGCTGAAAGAACAGAATAATCGTAGCGGATC<br>CGGTGGTGGTAGCGAAAATCTGTATTTTCAAGGTATGGTGA<br>GCAAAGGCGAAGAACTGTTTACCGGTGTTGTTCCGATTCTG<br>GTTGAACTGGATGGTGACGTTAATGGTCACAAATTTTCAGT<br>TAGCGGTGAAGGCGAAGGTGATGCAACCTATGGTAAACTG<br>ACCCTGAAATTTATCTGTACCACCGGCAAACTGCCGGTGCC<br>GTGGCCGACACTGGTTACCACACTGACCTATGGTGTTCAGT<br>GTTTTAGCCGTTATCCGGATCACATGAAACAGCACGATTTC<br>TTTAAAAGCGCAATGCCGGAAGGTTATGTTCAAGAACGTAC<br>CATCTTCTTTAAGGATGACGGCAACTATAAAACCCGTGCCG<br>AAGTTAAATTTGAAGGTGATACCCTGGTGAATCGCATTGAA<br>CTGAAAGGCATTGATTTCAAAGAGGATGGTAATATCCTGGG<br>CCACAAACTGGAATATAATTATAATAGCCACAACGTGTACAT<br>CATGGCCGACAAACAGAAAAATGGCATCAAAGTGAACTTCA<br>AGATCCGCCATAATATTGAAGATGGTTCAGTTCAGCTGGCC<br>GATCATTATCAGCAGAATACCCCGATCGGTGATGGTCCTGT<br>TCTGCTGCCTGATAATCATTATCTGAGCACCCAGAGCAAAC<br>TGAGCAAAGATCCGAATGAAAAACGTGATCACATGGTCCTG<br>CTGGAATTTGTTACCGCAGCAGGTATTACCTTAGGTATGGA<br>TGAACTGTACAAATAA |

**Table S2.** Sequences of gBlocks used in Figure 3. All sequences were ordered with a 5' T7 Promoter and ribosome binding site and a 3' T7 Terminator.

| Construct | DNA Sequence |
|---|---|
| MscL-GFP (MSIIK) | TAATACGACTCACTATAGGGGAATTGTGAGCGGATAAC<br>AATTCCCCTCTAGAAATAATTTTGTTTAACTTTAAGAAGG<br>AGATATACCATGGGCATGAGCATTATCAAAGAATTTCG<br>CGAATTTGCGATGCGCGGGAACGTGGTGGATTTGGCG<br>GTGGGTGTCATTATCGGTGCGGCATTCGGGAAGATTGT<br>CTCTTCACTGGTTGCCGATATCATCATGCCTCCTCTGG<br>GCTTATTAATTGGCGGGATCGATTTTAAACAGTTTGCTG<br>TCACGCTACGCGATGCGCAGGGGGATATCCCTGCTGT<br>TGTGATGCATTACGGTGTCTTCATTCAAAACGTCTTTGA<br>TTTTCTGATTGTGGCCTTTGCCATCTTTATGGCGATTAA<br>GCTAATCAACAAACTGAATCGGAAAAAAGAAGAACCAG<br>CAGCCGCACCTGCACCAACTAAAGAAGAAGTATTACTG<br>ACAGAAATTCGTGATTTGCTGAAAGAGCAGAATAACCG<br>CTCTGGATCCGGCGGCGGCAGCGAAAACCTGTATTTTC<br>AGGGCATGGTGAGCAAGGGCGAGGAGCTGTTCACCGG<br>GGTGGTGCCCATCCTGGTCGAGCTGGACGGCGACGTA<br>AACGGCCACAAGTTCAGCGTGTCCGGCGAGGGCGAGG<br>GCGATGCCACCTACGGCAAGCTGACCCTGAAGTTCATC<br>TGCACCACCGGCAAGCTGCCCGTGCCCTGGCCCACCC<br>TCGTGACCACCCTGACCTACGGCGTGCAGTGCTTCAG<br>CCGCTACCCCGACCACATGAAGCAGCACGACTTCTTCA<br>AGTCCGCCATGCCCGAAGGCTACGTCCAGGAGCGCAC<br>CATCTTCTTCAAGGACGACGGCAACTACAAGACCCGCG<br>CCGAGGTGAAGTTCGAGGGCGACACCCTGGTGAACCG<br>CATCGAGCTGAAGGGCATCGACTTCAAGGAGGACGGC<br>AACATCCTGGGGCACAAGCTGGAGTACAACTACAACAG<br>CCACAACGTCTATATCATGGCCGACAAGCAGAAGAACG<br>GCATCAAGGTGAACTTCAAGATCCGCCACAACATCGAG<br>GACGGCAGCGTGCAGCTCGCCGACCACTACCAGCAGA<br>ACACCCCCATCGGCGACGGCCCCGTGCTGCTGCCCGA<br>CAACCACTACCTGAGCACCCAGTCCAAGCTGAGCAAAG<br>ACCCCAACGAGAAGCGCGATCACATGGTCCTGCTGGA<br>GTTCGTGACCGCCGCCGGGATCACTCTCGGCATGGAC<br>GAGCTGTACAAGTAAGGATCCGGCTGCTAACAAAGCCC<br>GAAAGGAAGCTGAGTTGGCTGCTGCCACCGCTGAGCA<br>ATAACTAGCATAACCCCTTGGGGCCTCTAAACGGGTCT<br>TGAGGGGTTTTTTG |
| MAFFK-MscL-GFP | TAATACGACTCACTATAGGGGAATTGTGAGCGGATAAC<br>AATTCCCCTCTAGAAATAATTTTGTTTAACTTTAAGAAGG<br>AGATATACCATGGGCATGGCGTTTTTCAAAGAATTTCGC<br>GAATTTGCGATGCGCGGGAACGTGGTGGATTTGGCGG<br>TGGGTGTCATTATCGGTGCGGCATTCGGGAAGATTGTC<br>TCTTCACTGGTTGCCGATATCATCATGCCTCCTCTGGG<br>CTTATTAATTGGCGGGATCGATTTTAAACAGTTTGCTGT<br>CACGCTACGCGATGCGCAGGGGGATATCCCTGCTGTT<br>GTGATGCATTACGGTGTCTTCATTCAAAACGTCTTTGAT<br>TTTCTGATTGTGGCCTTTGCCATCTTTATGGCGATTAAG<br>CTAATCAACAAACTGAATCGGAAAAAAGAAGAACCAGC<br>AGCCGCACCTGCACCAACTAAAGAAGAAGTATTACTGA<br>CAGAAATTCGTGATTTGCTGAAAGAGCAGAATAACCGC<br>TCTGGATCCGGCGGCGGCAGCGAAAACCTGTATTTTCA |

| | |
|---|---|
| | GGGCATGGTGAGCAAGGGCGAGGAGCTGTTCACCGG<br>GGTGGTGCCCATCCTGGTCGAGCTGGACGGCGACGTA<br>AACGGCCACAAGTTCAGCGTGTCCGGCGAGGGCGAGG<br>GCGATGCCACCTACGGCAAGCTGACCCTGAAGTTCATC<br>TGCACCACCGGCAAGCTGCCCGTGCCCTGGCCCACCC<br>TCGTGACCACCCTGACCTACGGCGTGCAGTGCTTCAG<br>CCGCTACCCCGACCACATGAAGCAGCACGACTTCTTCA<br>AGTCCGCCATGCCCGAAGGCTACGTCCAGGAGCGCAC<br>CATCTTCTTCAAGGACGACGGCAACTACAAGACCCGCG<br>CCGAGGTGAAGTTCGAGGGCGACACCCTGGTGAACCG<br>CATCGAGCTGAAGGGCATCGACTTCAAGGAGGACGGC<br>AACATCCTGGGGCACAAGCTGGAGTACAACTACAACAG<br>CCACAACGTCTATATCATGGCCGACAAGCAGAAGAACG<br>GCATCAAGGTGAACTTCAAGATCCGCCACAACATCGAG<br>GACGGCAGCGTGCAGCTCGCCGACCACTACCAGCAGA<br>ACACCCCCATCGGCGACGGCCCCGTGCTGCTGCCCGA<br>CAACCACTACCTGAGCACCCAGTCCAAGCTGAGCAAAG<br>ACCCCAACGAGAAGCGCGATCACATGGTCCTGCTGGA<br>GTTCGTGACCGCCGCCGGGATCACTCTCGGCATGGAC<br>GAGCTGTACAAGTAAGGATCCGGCTGCTAACAAAGCCC<br>GAAAGGAAGCTGAGTTGGCTGCTGCCACCGCTGAGCA<br>ATAACTAGCATAACCCCTTGGGGCCTCTAAACGGGTCT<br>TGAGGGGTTTTTTG |
| MGIAK-MscL-GFP | TAATACGACTCACTATAGGGGAATTGTGAGCGGATAAC<br>AATTCCCCTCTAGAAATAATTTTGTTTAACTTTAAGAAGG<br>AGATATACCATGGGCATGGGCATTGCGAAAGAATTTCG<br>CGAATTTGCGATGCGCGGGAACGTGGTGGATTTGGCG<br>GTGGGTGTCATTATCGGTGCGGCATTCGGGAAGATTGT<br>CTCTTCACTGGTTGCCGATATCATCATGCCTCCTCTGG<br>GCTTATTAATTGGCGGGATCGATTTTAAACAGTTTGCTG<br>TCACGCTACGCGATGCGCAGGGGGATATCCCTGCTGT<br>TGTGATGCATTACGGTGTCTTCATTCAAAACGTCTTTGA<br>TTTTCTGATTGTGGCCTTTGCCATCTTTATGGCGATTAA<br>GCTAATCAACAAACTGAATCGGAAAAAAGAAGAACCAG<br>CAGCCGCACCTGCACCAACTAAAGAAGAAGTATTACTG<br>ACAGAAATTCGTGATTTGCTGAAAGAGCAGAATAACCG<br>CTCTGGATCCGGCGGCGGCAGCGAAAACCTGTATTTTC<br>AGGGCATGGTGAGCAAGGGCGAGGAGCTGTTCACCGG<br>GGTGGTGCCCATCCTGGTCGAGCTGGACGGCGACGTA<br>AACGGCCACAAGTTCAGCGTGTCCGGCGAGGGCGAGG<br>GCGATGCCACCTACGGCAAGCTGACCCTGAAGTTCATC<br>TGCACCACCGGCAAGCTGCCCGTGCCCTGGCCCACCC<br>TCGTGACCACCCTGACCTACGGCGTGCAGTGCTTCAG<br>CCGCTACCCCGACCACATGAAGCAGCACGACTTCTTCA<br>AGTCCGCCATGCCCGAAGGCTACGTCCAGGAGCGCAC<br>CATCTTCTTCAAGGACGACGGCAACTACAAGACCCGCG<br>CCGAGGTGAAGTTCGAGGGCGACACCCTGGTGAACCG<br>CATCGAGCTGAAGGGCATCGACTTCAAGGAGGACGGC<br>AACATCCTGGGGCACAAGCTGGAGTACAACTACAACAG<br>CCACAACGTCTATATCATGGCCGACAAGCAGAAGAACG<br>GCATCAAGGTGAACTTCAAGATCCGCCACAACATCGAG<br>GACGGCAGCGTGCAGCTCGCCGACCACTACCAGCAGA<br>ACACCCCCATCGGCGACGGCCCCGTGCTGCTGCCCGA<br>CAACCACTACCTGAGCACCCAGTCCAAGCTGAGCAAAG<br>ACCCCAACGAGAAGCGCGATCACATGGTCCTGCTGGA<br>GTTCGTGACCGCCGCCGGGATCACTCTCGGCATGGAC |

| | |
|---|---|
| | GAGCTGTACAAGTAAGGATCCGGCTGCTAACAAAGCCC<br>GAAAGGAAGCTGAGTTGGCTGCTGCCACCGCTGAGCA<br>ATAACTAGCATAACCCCTTGGGGCCTCTAAACGGGTCT<br>TGAGGGGTTTTTTG |
| MIKEF-MscL-GFP | TAATACGACTCACTATAGGGGAATTGTGAGCGGATAAC<br>AATTCCCCTCTAGAAATAATTTTGTTTAACTTTAAGAAGG<br>AGATATACCATGGGCATGATTAAAGAATTTGAATTTCGC<br>GAATTTGCGATGCGCGGGAACGTGGTGGATTTGGCGG<br>TGGGTGTCATTATCGGTGCGGCATTCGGGAAGATTGTC<br>TCTTCACTGGTTGCCGATATCATCATGCCTCCTCTGGG<br>CTTATTAATTGGCGGGATCGATTTTAAACAGTTTGCTGT<br>CACGCTACGCGATGCGCAGGGGGATATCCCTGCTGTT<br>GTGATGCATTACGGTGTCTTCATTCAAAACGTCTTTGAT<br>TTTCTGATTGTGGCCTTTGCCATCTTTATGGCGATTAAG<br>CTAATCAACAAACTGAATCGGAAAAAAGAAGAACCAGC<br>AGCCGCACCTGCACCAACTAAAGAAGAAGTATTACTGA<br>CAGAAATTCGTGATTTGCTGAAAGAGCAGAATAACCGC<br>TCTGGATCCGGCGGCGGCAGCGAAAACCTGTATTTTCA<br>GGGCATGGTGAGCAAGGGCGAGGAGCTGTTCACCGG<br>GGTGGTGCCCATCCTGGTCGAGCTGGACGGCGACGTA<br>AACGGCCACAAGTTCAGCGTGTCCGGCGAGGGCGAGG<br>GCGATGCCACCTACGGCAAGCTGACCCTGAAGTTCATC<br>TGCACCACCGGCAAGCTGCCCGTGCCCTGGCCCACCC<br>TCGTGACCACCCTGACCTACGGCGTGCAGTGCTTCAG<br>CCGCTACCCCGACCACATGAAGCAGCACGACTTCTTCA<br>AGTCCGCCATGCCCGAAGGCTACGTCCAGGAGCGCAC<br>CATCTTCTTCAAGGACGACGGCAACTACAAGACCCGCG<br>CCGAGGTGAAGTTCGAGGGCGACACCCTGGTGAACCG<br>CATCGAGCTGAAGGGCATCGACTTCAAGGAGGACGGC<br>AACATCCTGGGGCACAAGCTGGAGTACAACTACAACAG<br>CCACAACGTCTATATCATGGCCGACAAGCAGAAGAACG<br>GCATCAAGGTGAACTTCAAGATCCGCCACAACATCGAG<br>GACGGCAGCGTGCAGCTCGCCGACCACTACCAGCAGA<br>ACACCCCCATCGGCGACGGCCCCGTGCTGCTGCCCGA<br>CAACCACTACCTGAGCACCCAGTCCAAGCTGAGCAAAG<br>ACCCCAACGAGAAGCGCGATCACATGGTCCTGCTGGA<br>GTTCGTGACCGCCGCCGGGATCACTCTCGGCATGGAC<br>GAGCTGTACAAGTAAGGATCCGGCTGCTAACAAAGCCC<br>GAAAGGAAGCTGAGTTGGCTGCTGCCACCGCTGAGCA<br>ATAACTAGCATAACCCCTTGGGGCCTCTAAACGGGTCT<br>TGAGGGGTTTTTTG |
| MYYLK-MscL-GFP | TAATACGACTCACTATAGGGGAATTGTGAGCGGATAAC<br>AATTCCCCTCTAGAAATAATTTTGTTTAACTTTAAGAAGG<br>AGATATACCATGGGCATGTATTACCTGAAAGAATTTCGC<br>GAATTTGCGATGCGCGGGAACGTGGTGGATTTGGCGG<br>TGGGTGTCATTATCGGTGCGGCATTCGGGAAGATTGTC<br>TCTTCACTGGTTGCCGATATCATCATGCCTCCTCTGGG<br>CTTATTAATTGGCGGGATCGATTTTAAACAGTTTGCTGT<br>CACGCTACGCGATGCGCAGGGGGATATCCCTGCTGTT<br>GTGATGCATTACGGTGTCTTCATTCAAAACGTCTTTGAT<br>TTTCTGATTGTGGCCTTTGCCATCTTTATGGCGATTAAG<br>CTAATCAACAAACTGAATCGGAAAAAAGAAGAACCAGC<br>AGCCGCACCTGCACCAACTAAAGAAGAAGTATTACTGA<br>CAGAAATTCGTGATTTGCTGAAAGAGCAGAATAACCGC<br>TCTGGATCCGGCGGCGGCAGCGAAAACCTGTATTTTCA<br>GGGCATGGTGAGCAAGGGCGAGGAGCTGTTCACCGG |

| | |
|---|---|
| | GGTGGTGCCCATCCTGGTCGAGCTGGACGGCGACGTA<br>AACGGCCACAAGTTCAGCGTGTCCGGCGAGGGCGAGG<br>GCGATGCCACCTACGGCAAGCTGACCCTGAAGTTCATC<br>TGCACCACCGGCAAGCTGCCCGTGCCCTGGCCCACCC<br>TCGTGACCACCCTGACCTACGGCGTGCAGTGCTTCAG<br>CCGCTACCCCGACCACATGAAGCAGCACGACTTCTTCA<br>AGTCCGCCATGCCCGAAGGCTACGTCCAGGAGCGCAC<br>CATCTTCTTCAAGGACGACGGCAACTACAAGACCCGCG<br>CCGAGGTGAAGTTCGAGGGCGACACCCTGGTGAACCG<br>CATCGAGCTGAAGGGCATCGACTTCAAGGAGGACGGC<br>AACATCCTGGGGCACAAGCTGGAGTACAACTACAACAG<br>CCACAACGTCTATATCATGGCCGACAAGCAGAAGAACG<br>GCATCAAGGTGAACTTCAAGATCCGCCACAACATCGAG<br>GACGGCAGCGTGCAGCTCGCCGACCACTACCAGCAGA<br>ACACCCCCATCGGCGACGGCCCCGTGCTGCTGCCCGA<br>CAACCACTACCTGAGCACCCAGTCCAAGCTGAGCAAAG<br>ACCCCAACGAGAAGCGCGATCACATGGTCCTGCTGGA<br>GTTCGTGACCGCCGCCGGGATCACTCTCGGCATGGAC<br>GAGCTGTACAAGTAAGGATCCGGCTGCTAACAAAGCCC<br>GAAAGGAAGCTGAGTTGGCTGCTGCCACCGCTGAGCA<br>ATAACTAGCATAACCCCTTGGGGCCTCTAAACGGGTCT<br>TGAGGGGTTTTTTG |
| Polyleucine – MscL- GFP | TAATACGACTCACTATAGGGGAATTGTGAGCGGATAAC<br>AATTCCCCTCTAGAAATAATTTTGTTTAACTTTAAGAAGG<br>AGATATACCATGGGCATGCTGTTACTTCTGGAATTTCGC<br>GAATTTGCGATGCGCGGGAACGTGGTGGATTTGGCGG<br>TGGGTGTCATTATCGGTGCGGCATTCGGGAAGATTGTC<br>TCTTCACTGGTTGCCGATATCATCATGCCTCCTCTGGG<br>CTTATTAATTGGCGGGATCGATTTTAAACAGTTTGCTGT<br>CACGCTACGCGATGCGCAGGGGGATATCCCTGCTGTT<br>GTGATGCATTACGGTGTCTTCATTCAAAACGTCTTTGAT<br>TTTCTGATTGTGGCCTTTGCCATCTTTATGGCGATTAAG<br>CTAATCAACAAACTGAATCGGAAAAAAGAAGAACCAGC<br>AGCCGCACCTGCACCAACTAAAGAAGAAGTATTACTGA<br>CAGAAATTCGTGATTTGCTGAAAGAGCAGAATAACCGC<br>TCTGGATCCGGCGGCGGCAGCGAAAACCTGTATTTTCA<br>GGGCATGGTGAGCAAGGGCGAGGAGCTGTTCACCGG<br>GGTGGTGCCCATCCTGGTCGAGCTGGACGGCGACGTA<br>AACGGCCACAAGTTCAGCGTGTCCGGCGAGGGCGAGG<br>GCGATGCCACCTACGGCAAGCTGACCCTGAAGTTCATC<br>TGCACCACCGGCAAGCTGCCCGTGCCCTGGCCCACCC<br>TCGTGACCACCCTGACCTACGGCGTGCAGTGCTTCAG<br>CCGCTACCCCGACCACATGAAGCAGCACGACTTCTTCA<br>AGTCCGCCATGCCCGAAGGCTACGTCCAGGAGCGCAC<br>CATCTTCTTCAAGGACGACGGCAACTACAAGACCCGCG<br>CCGAGGTGAAGTTCGAGGGCGACACCCTGGTGAACCG<br>CATCGAGCTGAAGGGCATCGACTTCAAGGAGGACGGC<br>AACATCCTGGGGCACAAGCTGGAGTACAACTACAACAG<br>CCACAACGTCTATATCATGGCCGACAAGCAGAAGAACG<br>GCATCAAGGTGAACTTCAAGATCCGCCACAACATCGAG<br>GACGGCAGCGTGCAGCTCGCCGACCACTACCAGCAGA<br>ACACCCCCATCGGCGACGGCCCCGTGCTGCTGCCCGA<br>CAACCACTACCTGAGCACCCAGTCCAAGCTGAGCAAAG<br>ACCCCAACGAGAAGCGCGATCACATGGTCCTGCTGGA<br>GTTCGTGACCGCCGCCGGGATCACTCTCGGCATGGAC<br>GAGCTGTACAAGTAAGGATCCGGCTGCTAACAAAGCCC |

| | |
|---|---|
| | GAAAGGAAGCTGAGTTGGCTGCTGCCACCGCTGAGCA<br>ATAACTAGCATAACCCCTTGGGGCCTCTAAACGGGTCT<br>TGAGGGGTTTTTTG |
| 20 Å Protein | TAATACGACTCACTATAGGGGAATTGTGAGCGGATAAC<br>AATTCCCCTCTAGAAATAATTTTGTTTAACTTTAAGAAGG<br>AGATATACCATGGGTAGCACCCGCAAAGAAATTATCGA<br>AAAACTGGAAAAATCACTGCGTCGCCAGAAAAAGCTGG<br>CACGTTTTCTGCTGATTCTGCTGCTGCTGTTACTGGCA<br>CTGCTGCTGGAACTGTTAGAACTGCTGCGTCGTCTGGA<br>AGAACTGCAACGTCGTGGTAGCAGTGATGAAGAAGTG<br>CACGAACTGTTACGTCGTATTATTGAACTGGTGGAATAT<br>ATCATCCTGCTGGTCCTGTTTATTATCGTTCTGGTGCGC<br>ATTATTATCAAACTGGCAGAACATCAGCGTCGCCTGGT<br>TGAAGAACTGAAAAAGCAGGATGGATCCGGCGGCGGC<br>AGCGAAAACCTGTATTTTCAGGGCATGGTGAGCAAGGG<br>CGAGGAGCTGTTCACCGGGGTGGTGCCCATCCTGGTC<br>GAGCTGGACGGCGACGTAAACGGCCACAAGTTCAGCG<br>TGTCCGGCGAGGGCGAGGGCGATGCCACCTACGGCA<br>AGCTGACCCTGAAGTTCATCTGCACCACCGGCAAGCTG<br>CCCGTGCCCTGGCCCACCCTCGTGACCACCCTGACCT<br>ACGGCGTGCAGTGCTTCAGCCGCTACCCCGACCACAT<br>GAAGCAGCACGACTTCTTCAAGTCCGCCATGCCCGAA<br>GGCTACGTCCAGGAGCGCACCATCTTCTTCAAGGACG<br>ACGGCAACTACAAGACCCGCGCCGAGGTGAAGTTCGA<br>GGGCGACACCCTGGTGAACCGCATCGAGCTGAAGGGC<br>ATCGACTTCAAGGAGGACGGCAACATCCTGGGGCACA<br>AGCTGGAGTACAACTACAACAGCCACAACGTCTATATC<br>ATGGCCGACAAGCAGAAGAACGGCATCAAGGTGAACT<br>TCAAGATCCGCCACAACATCGAGGACGGCAGCGTGCA<br>GCTCGCCGACCACTACCAGCAGAACACCCCCATCGGC<br>GACGGCCCCGTGCTGCTGCCCGACAACCACTACCTGA<br>GCACCCAGTCCAAGCTGAGCAAAGACCCCAACGAGAA<br>GCGCGATCACATGGTCCTGCTGGAGTTCGTGACCGCC<br>GCCGGGATCACTCTCGGCATGGACGAGCTGTACAAGT<br>AAGGATCCGGCTGCTAACAAAGCCCGAAAGGAAGCTG<br>AGTTGGCTGCTGCCACCGCTGAGCAATAACTAGCATAA<br>CCCCTTGGGGCCTCTAAACGGGTCTTGAGGGGTTTTT<br>G |
| 20 Å Protein (G2Y) | TAATACGACTCACTATAGGGGAATTGTGAGCGGATAAC<br>AATTCCCCTCTAGAAATAATTTTGTTTAACTTTAAGAAGG<br>AGATATACCATGGGTAGCACCCGCAAATTCATTATCTTC<br>AAACTGGAAAAATCACTGCGTCGCCAGAAAAAGCTGGC<br>ACGTTTTCTGCTGATTCTGCTGCTGCTGTTACTGGCACT<br>GCTGCTGGAACTGTTAGAACTGCTGCGTCGTCTGGAAG<br>AACTGCAACGTCGTGGTAGCAGTGATGAAGAAGTGCAC<br>GAACTGTTACGTCGTATTATTGAACTGGTGGAATATATC<br>ATCCTGCTGGTCCTGTTTATTATCGTTCTGGTGCGCATT<br>ATTATCAAACTGGCAGAACATCAGCGTCGCCTGGTTGA<br>AGAACTGAAAAAGCAGGATGGATCCGGCGGCGGCAGC<br>GAAAACCTGTATTTTCAGGGCATGGTGAGCAAGGGCGA<br>GGAGCTGTTCACCGGGGTGGTGCCCATCCTGGTCGAG<br>CTGGACGGCGACGTAAACGGCCACAAGTTCAGCGTGT<br>CCGGCGAGGGCGAGGGCGATGCCACCTACGGCAAGC<br>TGACCCTGAAGTTCATCTGCACCACCGGCAAGCTGCCC<br>GTGCCCTGGCCCACCCTCGTGACCACCCTGACCTACG |

| | |
|---|---|
| | GCGTGCAGTGCTTCAGCCGCTACCCCGACCACATGAA<br>GCAGCACGACTTCTTCAAGTCCGCCATGCCCGAAGGCT<br>ACGTCCAGGAGCGCACCATCTTCTTCAAGGACGACGG<br>CAACTACAAGACCCGCGCCGAGGTGAAGTTCGAGGGC<br>GACACCCTGGTGAACCGCATCGAGCTGAAGGGCATCG<br>ACTTCAAGGAGGACGGCAACATCCTGGGGCACAAGCT<br>GGAGTACAACTACAACAGCCACAACGTCTATATCATGG<br>CCGACAAGCAGAAGAACGGCATCAAGGTGAACTTCAA<br>GATCCGCCACAACATCGAGGACGGCAGCGTGCAGCTC<br>GCCGACCACTACCAGCAGAACACCCCCATCGGCGACG<br>GCCCCGTGCTGCTGCCCGACAACCACTACCTGAGCAC<br>CCAGTCCAAGCTGAGCAAAGACCCCAACGAGAAGCGC<br>GATCACATGGTCCTGCTGGAGTTCGTGACCGCCGCCG<br>GGATCACTCTCGGCATGGACGAGCTGTACAAGTAAGG<br>ATCCGGCTGCTAACAAAGCCCGAAAGGAAGCTGAGTT<br>GGCTGCTGCCACCGCTGAGCAATAACTAGCATAACCCC<br>TTGGGGCCTCTAAACGGGTCTTGAGGGGTTTTTTG |
| 20 Å Protein (G2Y, S3Y) | TAATACGACTCACTATAGGGGAATTGTGAGCGGATAAC<br>AATTCCCCTCTAGAAATAATTTTGTTTAACTTTAAGAAGG<br>AGATATACCATGTACTACACCCGCAAAGAAATTATCGAA<br>AAACTGGAAAAATCACTGCGTCGCCAGAAAAAGCTGGC<br>ACGTTTTCTGCTGATTCTGCTGCTGCTGTTACTGGCACT<br>GCTGCTGGAACTGTTAGAACTGCTGCGTCGTCTGGAAG<br>AACTGCAACGTCGTGGTAGCAGTGATGAAGAAGTGCAC<br>GAACTGTTACGTCGTATTATTGAACTGGTGGAATATATC<br>ATCCTGCTGGTCCTGTTTATTATCGTTCTGGTGCGCATT<br>ATTATCAAACTGGCAGAACATCAGCGTCGCCTGGTTGA<br>AGAACTGAAAAAGCAGGATGGATCCGGCGGCGGCAGC<br>GAAAACCTGTATTTTCAGGGCATGGTGAGCAAGGGCGA<br>GGAGCTGTTCACCGGGGTGGTGCCCATCCTGGTCGAG<br>CTGGACGGCGACGTAAACGGCCACAAGTTCAGCGTGT<br>CCGGCGAGGGCGAGGGCGATGCCACCTACGGCAAGC<br>TGACCCTGAAGTTCATCTGCACCACCGGCAAGCTGCCC<br>GTGCCCTGGCCCACCCTCGTGACCACCCTGACCTACG<br>GCGTGCAGTGCTTCAGCCGCTACCCCGACCACATGAA<br>GCAGCACGACTTCTTCAAGTCCGCCATGCCCGAAGGCT<br>ACGTCCAGGAGCGCACCATCTTCTTCAAGGACGACGG<br>CAACTACAAGACCCGCGCCGAGGTGAAGTTCGAGGGC<br>GACACCCTGGTGAACCGCATCGAGCTGAAGGGCATCG<br>ACTTCAAGGAGGACGGCAACATCCTGGGGCACAAGCT<br>GGAGTACAACTACAACAGCCACAACGTCTATATCATGG<br>CCGACAAGCAGAAGAACGGCATCAAGGTGAACTTCAA<br>GATCCGCCACAACATCGAGGACGGCAGCGTGCAGCTC<br>GCCGACCACTACCAGCAGAACACCCCCATCGGCGACG<br>GCCCCGTGCTGCTGCCCGACAACCACTACCTGAGCAC<br>CCAGTCCAAGCTGAGCAAAGACCCCAACGAGAAGCGC<br>GATCACATGGTCCTGCTGGAGTTCGTGACCGCCGCCG<br>GGATCACTCTCGGCATGGACGAGCTGTACAAGTAAGG<br>ATCCGGCTGCTAACAAAGCCCGAAAGGAAGCTGAGTT<br>GGCTGCTGCCACCGCTGAGCAATAACTAGCATAACCCC<br>TTGGGGCCTCTAAACGGGTCTTGAGGGGTTTTTTG |
| 20 Å Protein (E7F, E10F) | TAATACGACTCACTATAGGGGAATTGTGAGCGGATAAC<br>AATTCCCCTCTAGAAATAATTTTGTTTAACTTTAAGAAGG<br>AGATATACCATGGGTAGCACCCGCAAATTCATTATCTTC |

| | |
|---|---|
| | AAACTGGAAAAATCACTGCGTCGCCAGAAAAAGCTGGC<br>ACGTTTTCTGCTGATTCTGCTGCTGCTGTTACTGGCACT<br>GCTGCTGGAACTGTTAGAACTGCTGCGTCGTCTGGAAG<br>AACTGCAACGTCGTGGTAGCAGTGATGAAGAAGTGCAC<br>GAACTGTTACGTCGTATTATTGAACTGGTGGAATATATC<br>ATCCTGCTGGTCCTGTTTATTATCGTTCTGGTGCGCATT<br>ATTATCAAACTGGCAGAACATCAGCGTCGCCTGGTTGA<br>AGAACTGAAAAAGCAGGATGGATCCGGCGGCGGCAGC<br>GAAAACCTGTATTTTCAGGGCATGGTGAGCAAGGGCGA<br>GGAGCTGTTCACCGGGGTGGTGCCCATCCTGGTCGAG<br>CTGGACGGCGACGTAAACGGCCACAAGTTCAGCGTGT<br>CCGGCGAGGGCGAGGGCGATGCCACCTACGGCAAGC<br>TGACCCTGAAGTTCATCTGCACCACCGGCAAGCTGCCC<br>GTGCCCTGGCCCACCCTCGTGACCACCCTGACCTACG<br>GCGTGCAGTGCTTCAGCCGCTACCCCGACCACATGAA<br>GCAGCACGACTTCTTCAAGTCCGCCATGCCCGAAGGCT<br>ACGTCCAGGAGCGCACCATCTTCTTCAAGGACGACGG<br>CAACTACAAGACCCGCGCCGAGGTGAAGTTCGAGGGC<br>GACACCCTGGTGAACCGCATCGAGCTGAAGGGCATCG<br>ACTTCAAGGAGGACGGCAACATCCTGGGGCACAAGCT<br>GGAGTACAACTACAACAGCCACAACGTCTATATCATGG<br>CCGACAAGCAGAAGAACGGCATCAAGGTGAACTTCAA<br>GATCCGCCACAACATCGAGGACGGCAGCGTGCAGCTC<br>GCCGACCACTACCAGCAGAACACCCCCATCGGCGACG<br>GCCCCGTGCTGCTGCCCGACAACCACTACCTGAGCAC<br>CCAGTCCAAGCTGAGCAAAGACCCCAACGAGAAGCGC<br>GATCACATGGTCCTGCTGGAGTTCGTGACCGCCGCCG<br>GGATCACTCTCGGCATGGACGAGCTGTACAAGTAAGG<br>ATCCGGCTGCTAACAAAGCCCGAAAGGAAGCTGAGTT<br>GGCTGCTGCCACCGCTGAGCAATAACTAGCATAACCCC<br>TTGGGGCCTCTAAACGGGTCTTGAGGGGTTTTTTG |
| 20 Å Protein (E7L, E10L) | TAATACGACTCACTATAGGGGAATTGTGAGCGGATAAC<br>AATTCCCCTCTAGAAATAATTTTGTTTAACTTTAAGAAGG<br>AGATATACCATGGGTAGCACCCGCAAACTGATTATCCT<br>GAAACTGGAAAAATCACTGCGTCGCCAGAAAAAGCTGG<br>CACGTTTTCTGCTGATTCTGCTGCTGCTGTTACTGGCA<br>CTGCTGCTGGAACTGTTAGAACTGCTGCGTCGTCTGGA<br>AGAACTGCAACGTCGTGGTAGCAGTGATGAAGAAGTG<br>CACGAACTGTTACGTCGTATTATTGAACTGGTGGAATAT<br>ATCATCCTGCTGGTCCTGTTTATTATCGTTCTGGTGCGC<br>ATTATTATCAAACTGGCAGAACATCAGCGTCGCCTGGT<br>TGAAGAACTGAAAAAGCAGGATGGATCCGGCGGCGGC<br>AGCGAAAACCTGTATTTTCAGGGCATGGTGAGCAAGGG<br>CGAGGAGCTGTTCACCGGGGTGGTGCCCATCCTGGTC<br>GAGCTGGACGGCGACGTAAACGGCCACAAGTTCAGCG<br>TGTCCGGCGAGGGCGAGGGCGATGCCACCTACGGCA<br>AGCTGACCCTGAAGTTCATCTGCACCACCGGCAAGCTG<br>CCCGTGCCCTGGCCCACCCTCGTGACCACCCTGACCT<br>ACGGCGTGCAGTGCTTCAGCCGCTACCCCGACCACAT<br>GAAGCAGCACGACTTCTTCAAGTCCGCCATGCCCGAA<br>GGCTACGTCCAGGAGCGCACCATCTTCTTCAAGGACG<br>ACGGCAACTACAAGACCCGCGCCGAGGTGAAGTTCGA<br>GGGCGACACCCTGGTGAACCGCATCGAGCTGAAGGGC<br>ATCGACTTCAAGGAGGACGGCAACATCCTGGGGCACA<br>AGCTGGAGTACAACTACAACAGCCACAACGTCTATATC |

| | |
|---|---|
| | ATGGCCGACAAGCAGAAGAACGGCATCAAGGTGAACT<br>TCAAGATCCGCCACAACATCGAGGACGGCAGCGTGCA<br>GCTCGCCGACCACTACCAGCAGAACACCCCCATCGGC<br>GACGGCCCCGTGCTGCTGCCCGACAACCACTACCTGA<br>GCACCCAGTCCAAGCTGAGCAAAGACCCCAACGAGAA<br>GCGCGATCACATGGTCCTGCTGGAGTTCGTGACCGCC<br>GCCGGGATCACTCTCGGCATGGACGAGCTGTACAAGT<br>AAGGATCCGGCTGCTAACAAAGCCCGAAAGGAAGCTG<br>AGTTGGCTGCTGCCACCGCTGAGCAATAACTAGCATAA<br>CCCCTTGGGGCCTCTAAACGGGTCTTGAGGGGTTTTTT<br>G |
